# Clear cell renal cell carcinoma consensus transcriptomic programs reveal converging trajectories towards aggressive disease

**DOI:** 10.64898/2026.06.24.734364

**Authors:** Roy Elias, Vivek Nimgaonkar, Bingrui E. Xie, Yuqi Zhang, Archana Balan, Kathleen Noller, Nirmish Singla, Yasser Ged, Ezra Baraban, Genevieve L. Stein-O’Brien, Payal Kapur, James Brugarolas, Michael F. Ochs, Elana J. Fertig, Atul Deshpande, Srinivasan Yegnasubramanian

## Abstract

Clear cell renal cell carcinoma (ccRCC) is characterized by a branching genomic trajectory in which early biallelic *VHL* inactivation splits into *PBRM1*- and *BAP1*-mutant lineages. However, driver mutations alone do not account for the molecular and phenotypic heterogeneity. To dissect this heterogeneity, we developed a non-negative matrix factorization (NMF)-based gene expression analytical framework for identifying recurrent transcriptional programs across factorization dimensionalities and across datasets. We applied it to three curated datasets (IMmotion151, n = 823; JAVELIN Renal 101, n = 726; TCGA, n = 614) to define 17 consensus transcriptomic programs (CTPs). Mapping CTPs onto single-cell RNAseq (scRNAseq) of human ccRCC tumors and patient-derived tumorgraft models linked these programs to their cellular sources, distinguishing RCC-intrinsic, tumor-cell-extrinsic, and mixed programs. RCC-intrinsic CTPs associated with canonical drivers, including *VHL* (R1), *PBRM1* (R2), *BAP1* (R4), *PTEN*/*TSC1* (R3), *TFE3/TFEB* fusions (R5), NF2 (R6), and *CDKN2A*/*TP53* (MP-Prolif). Additional CTPs captured tumor microenvironment (TME) composition (TME-Tcell, TME-Myelo, TME-Endo, TME-Stroma) and biological processes active across multiple cellular compartments, including proliferation, Y-chromosome–linked expression in male tumors, ciliary biology, and translation. Trajectory inference methods revealed *PBRM1*-like and *BAP1*-like branches that converged on a shared aggressive late transcriptomic stage (TS) associated with higher nuclear grade, additional driver alterations, myeloid/stromal infiltration, and poor clinical outcomes. Spatial transcriptomics (and multiregional sequencing) of paired conventional ccRCC and sarcomatoid regions linked TS advancement with morphological progression and clonal evolution. After adjusting for stage, grade, and *BAP1*/*PBRM1* status, TS remained independently prognostic. Furthermore, our data suggest that immune checkpoint inhibitor combinations are particularly beneficial for MP-Prolif and not R1 utilizing specimens. In summary, we present an atlas of recurring transcriptomic programs in RCC and an ontological framework bridging genotype, tumor-cell-intrinsic gene expression, and microenvironment remodeling, with implications for risk stratification and treatment selection in ccRCC.

## Introduction

Renal cell carcinoma (RCC) is among the ten most common malignancies and is characterized by striking clinical heterogeneity, ranging from indolent disease to rapidly progressive, lethal cancer^1^. Although immune checkpoint inhibitors (ICI) and vascular endothelial growth factor (VEGF) targeted therapies have improved survival, outcomes vary and results are unpredictable^2–6^. In clinical practice, treatment selection is largely empirical and biomarkers are lacking^1^. Consequently, many patients likely receive suboptimal therapy, either through exposure to unnecessary toxicity or through under-treatment of aggressive disease. The lack of biomarkers is particularly costly in the adjuvant setting, where a large fraction of patients are cured with surgery alone, while most that ultimately relapse are undertreated with current adjuvant regimens^7^.

Clear cell renal cell carcinoma (ccRCC), the most common histological subtype^8^, has been extensively characterized. The initiating event is typically biallelic inactivation of *VHL*, followed by mutations in either *PBRM1* or *BAP1*, with subsequent acquisition of additional mutations and copy-number events that influence histopathological aggressiveness and metastatic potential^9–12^. This branching genomic architecture is supported by the prevalence and mutual exclusivity of recurrent driver events in patient cohorts^9,13–15^ and is reinforced by multi-regional sequencing studies^11^ and genetically engineered mouse models^10^. However, the precise effects of these alterations on tumor behaviors such as metastatic progression, tumor microenvironment (TME) remodeling, and therapeutic response, remain incompletely understood. In particular, genetic alterations alone often provide an insufficient explanation for phenotypic heterogeneity. For example, roughly 40% of cases lack a detectable *BAP1* or *PBRM1* mutation^9,13,14^. Moreover, tumors with similar mutation status can exhibit substantial phenotypic variation; *PBRM1*-mutant tumors, for example, exhibit conflicting clinical characteristics and TME composition across studies^16–20^. This variability is not fully explained by the accumulation of additional driver mutations. Together, these observations suggest that factors beyond individual driver alterations contribute to ccRCC phenotypes and clinical behavior.

One approach to dissect phenotypes involves the identification of genes with coordinated expression across samples. Non-negative matrix factorization (NMF) has emerged as a mainstay for inferring such programs^21–23^. NMF concurrently learns gene signatures that define phenotypes and weights each program activity across samples. Program activity can then be related to sample level annotation (such as genotype)^24^. One limitation of NMF application to bulk RNA sequencing is admixing of tumor and TME signals. Programs learned by NMF can reflect true tumor-cell states, differences in immune or stromal infiltration, or a combination of both. NMF, like unsupervised matrix decomposition methods, requires the user to specify the number of factors (K) to learn. The choice of K influences which biological processes are recovered: low K can merge distinct processes into single factors, while high K can fragment coherent processes across multiple factors. This dependency can be partially mitigated by aggregating recurrent programs inferred across a range of factorization dimensionalities^22,25,26^.

Indeed, current ccRCC transcriptomic subtypes are often driven by microenvironmental composition and stratify tumors along axes such as “angiogenic” and “inflamed”^19,27^.

While these frameworks have provided important insights and may be linked with treatment response, they capture tumor-cell-intrinsic and -extrinsic signals jointly and assign tumors to discrete subtypes despite substantial intratumoral heterogeneity^28,29^. Resolving the tumor-cell-intrinsic and microenvironmental contributions to bulk transcriptional signal, and quantifying their utilization continuously rather than categorically, may extend these frameworks to better capture the biological diversity within ccRCC. Single-cell (sc)RNAseq can address these issues by attributing programs to specific cell types and separating malignant from non-malignant signals. However, scRNAseq cohorts remain orders of magnitude smaller than bulk cohorts, limiting statistical power to relate cell-resolved programs to patient-level outcomes such as survival, treatment response, and rare genomic alterations.

To address these limitations, we developed a multi-dimensional cross-cohort NMF workflow that defines reproducible ccRCC transcriptomic programs from bulk RNAseq. The workflow identifies recurring NMF factors, referred to as consensus transcriptomic programs (CTPs), across independent datasets and a range of factorization dimensionalities, using cross-cohort recurrence as a constraint that prioritizes biology over dataset-specific structure. Rather than assigning tumors to discrete subtypes, it quantifies CTP usage within each sample, allowing individual tumors to engage multiple programs simultaneously, potentially better accommodating intratumoral heterogeneity. We then map CTPs to their cellular sources using scRNAseq and patient-derived tumorgraft models, in which species-of-origin alignment isolates malignant from microenvironmental transcripts, separating tumor-cell-intrinsic from tumor-cell-extrinsic components.

We applied this framework across ccRCC transcriptomic datasets to define a CTP atlas and relate program usage to genetic alterations, microenvironmental composition, and clinicopathological attributes. Focusing on tumor-cell-intrinsic CTPs, we identified branching ccRCC trajectories that converge on an aggressive end-state, establishing a transcriptomic stage (TS) axis strongly associated with driver mutations, nuclear grade, TME remodeling, and clinical outcomes. The atlas is disseminated as the Renal Cell Carcinoma Consensus Transcriptomic Program (rC3TP) R package, enabling CTP scoring in new datasets.

## Results

### Inferring consensus transcriptomic programs in clear cell renal cell carcinoma

To obtain consensus transcriptomic programs (CTPs) from multiple ccRCC bulk RNAseq datasets, we (1) performed NMF using the CoGAPS algorithm^21,30^ across a range of factorization ranks (k) on each dataset independently, (2) identified recurrent programs across datasets via graph-based community detection, (3) consolidated each community into a weighted transcriptomic program using a modified Borda aggregation across factors and ranks, yielding CTPs, and (4) scored samples relative to a permuted null distribution to quantify CTP utilization (**Figure 1A, Methods**).

**Figure 1.**
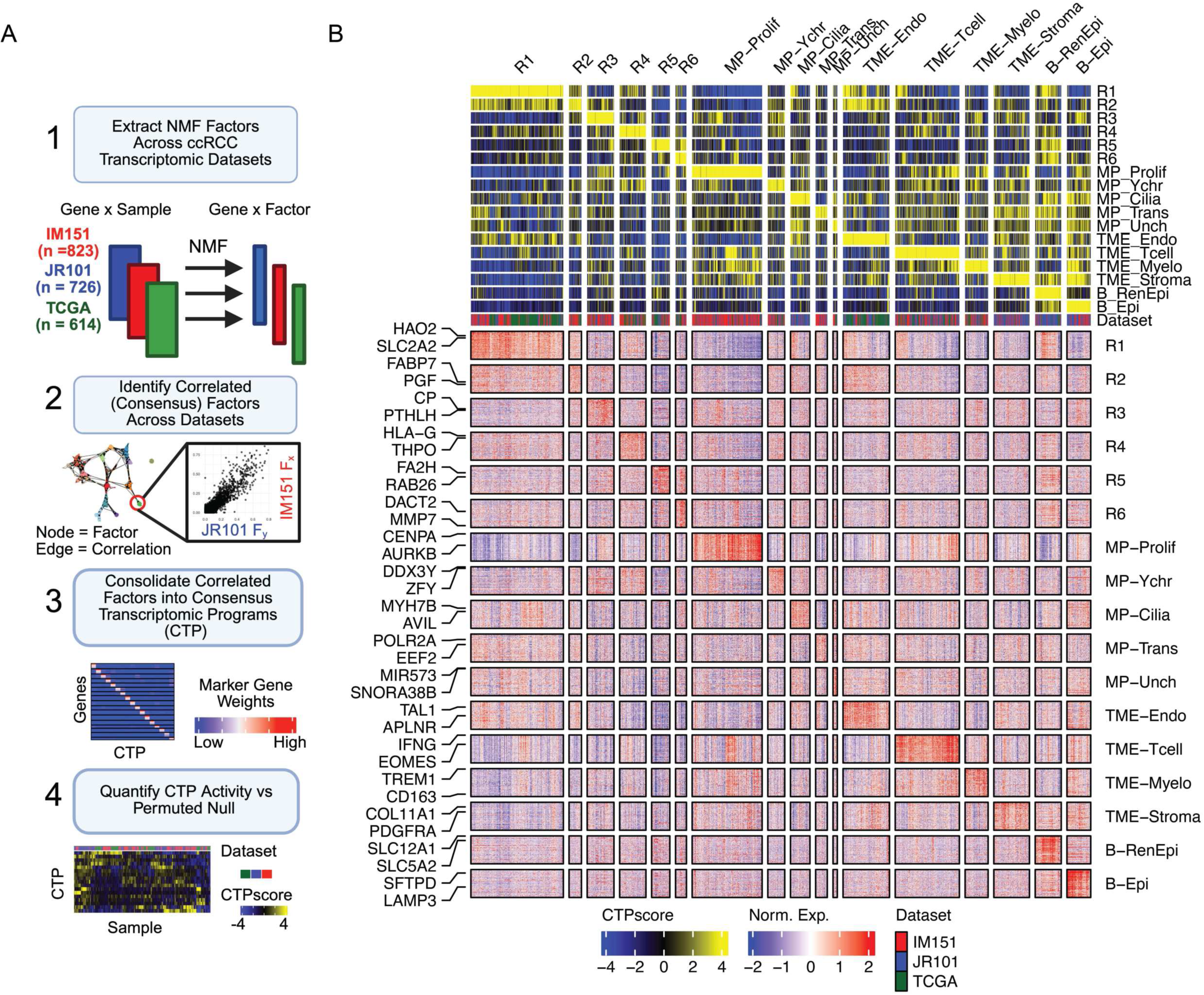
Defining consensus transcriptomic programs in clear cell renal cell carcinoma. (A) Schematic overview of the Consensus Transcriptomic Program (CTP) discovery workflow. Non-negative Matrix Factorization (NMF) was performed across a range of factorization (k) levels on three independent bulk RNAseq datasets (IM151^27^, JR101^5^, TCGA^9^). Inter-dataset correlated factors were identified using graph-based community detection and consolidated into weighted gene sets used to score individual samples (CTPscore; see **Methods**). (B) Heatmap of normalized gene expression for the top 100 marker genes defining each CTP. Samples (columns) are split by the maximally enriched CTP then arranged via unsupervised hierarchical clustering within each group. CTPscores and dataset of origin are displayed in the top annotation bars.

We applied this workflow to three ccRCC datasets comprising a total of 2,163 specimens: IMmotion151 (IM151^27^, n = 823), JAVELIN Renal 101^5^ (JR101, n = 726), and The Cancer Genome Atlas^9^ (TCGA, n = 614; **Table S1**). Across the three datasets and a range of factorization ranks, we obtained 840 NMF factors in total. Despite differences in sample processing (i.e., frozen vs formalin fixed), RNAseq platforms, and preprocessing pipelines, we observed high correlation of NMF factors learned from different datasets. Because the same algorithm and gene-selection procedure were applied to all three datasets, we asked whether this cross-dataset agreement could reflect a shared technical signature of the workflow rather than common biology. To test this, we performed the same NMF analysis on gene-wise permuted data, which preserves individual gene distributions while eliminating inter-gene correlations (**Figure S1A**). In the permuted data, factors remained dataset specific, suggesting that the cross-dataset correlations in the real data reflect biological signal rather than technical artifact (**Figure S1B**).

Given the high inter-dataset correlation among factors learned in the data, we hypothesized that a common set of features learned across values of (k) and between datasets would capture transcriptomic programs in ccRCC. To consolidate 840 factors into a representative set of gene expression programs, we first used a graph-based approach to plot factor pairs which exhibited a Pearson correlation ≥ 0.6, then identified communities of correlated factors using the Infomap community detection algorithm, which groups nodes that are more densely interconnected with each other than with the rest of the graph^31^. In the case of our bulk RNAseq cohort of ccRCC, this analysis resulted in 36 communities, of which 17 contained factors from at least two datasets with sufficient cross-dataset support (see Methods); we refer to these as consensus communities, encompassing 683 of the 840 (81%) input factors (**Figure S1C-D**). The remaining communities contained dataset-specific factors and were excluded. To enable per-sample scoring and downstream analysis, each consensus community was further consolidated into a weighted 100-gene signature using a Borda-ranking approach, yielding largely non-overlapping CTP marker gene sets (**Figure S1E, Table S2, see Methods**).

Having defined CTPs at the gene level, we next quantified their usage in individual samples. Standard transfer-learning approaches for projecting NMF factors between datasets were not applicable to our consolidated programs^32,33^. We instead computed CTP enrichment in each sample by summing, across the 100 marker genes, the product of each gene’s Borda-aggregated CTP weight (reflecting its relative importance within the program; **Methods**) and its log₂TPM expression. We then scaled these values against a null distribution derived by applying the same procedure to gene-wise permuted data, yielding a score that quantifies coordinated gene expression beyond chance and enables statistical testing; we refer to this as the CTPscore. CTPscores and CTP marker gene expression were concordant when visualized in a heatmap, supporting the interpretation that CTPs represent a collection of co-expressed genes (**Figure 1B**). More broadly, CTPscores were heterogeneous across the datasets, suggesting that they capture recurring phenotypes across datasets. Unlike clustering-based subtyping, this NMF-based framework allows for individual samples to be enriched for multiple CTPs simultaneously. We defined CTP utilization as a CTPscore significantly higher than the permuted null distribution (FDR < 0.05). CTP utilization frequencies ranged from 17-38% across CTPs, and the median number of CTPs utilized per sample was 4 (interquartile range [IQR] 3-6) (**Figure S2**).

### Consensus transcriptomic programs distinguish RCC-intrinsic from -extrinsic processes

We next sought to ascribe biological function to the CTPs. First, we annotated associations with existing gene-sets using a gene-set overrepresentation analysis^34^. Consistent with the non-overlapping marker genes comprising CTPs, CTPs were associated with non-overlapping gene sets. These gene sets captured diverse functional themes such as proliferation, immune regulation, solute transport, stromal remodeling, and Y-chromosome (chr) associated genes (**Figure 2A**). However, overrepresentation analysis provided only a partial view of CTP biology, with several programs lacking strong overlap with any established gene set. We therefore complemented this approach with orthogonal analyses to resolve the cellular sources and biological context of each CTP.

**Figure 2.**
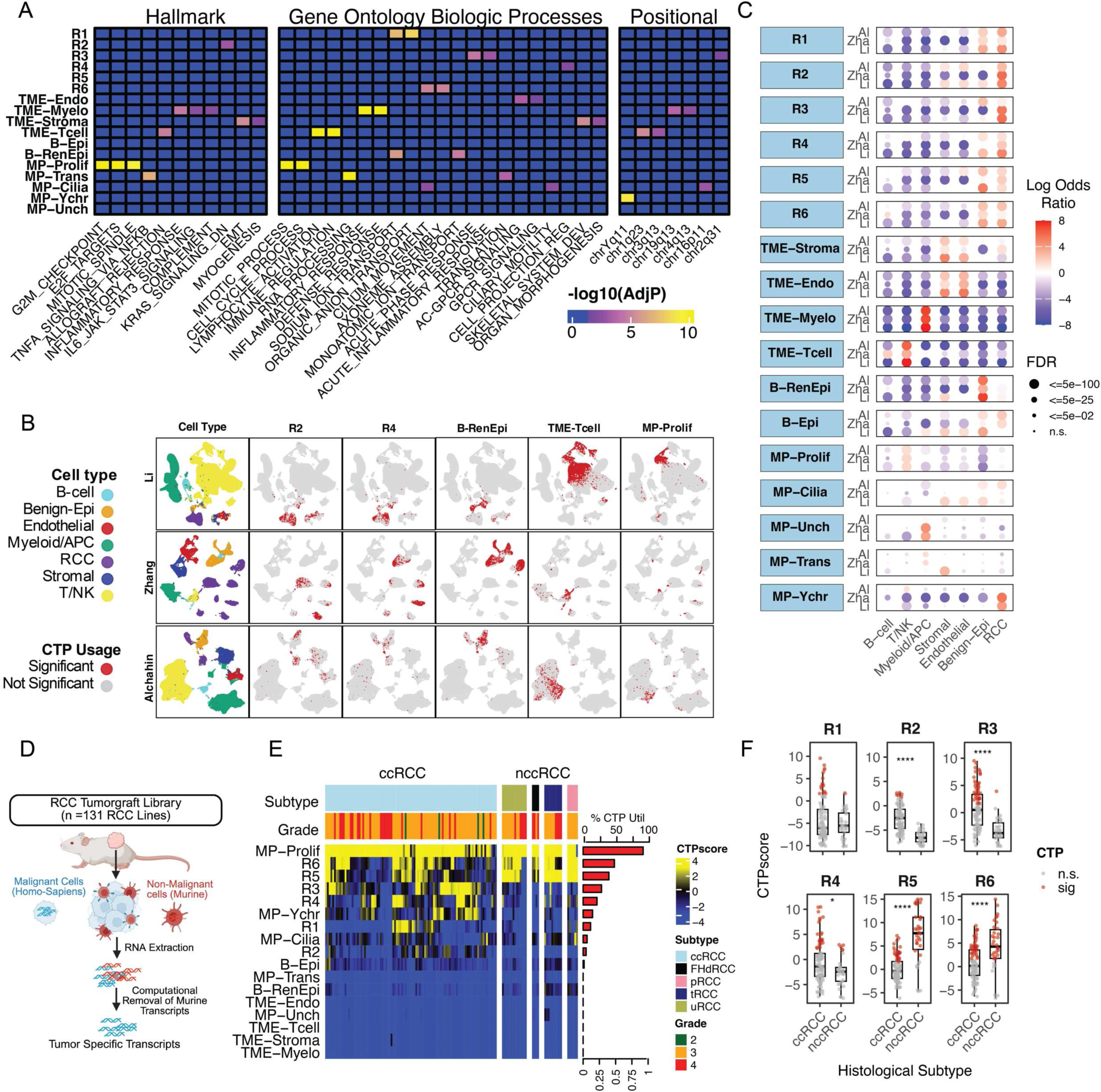
Mapping cellular source of RCC consensus transcriptomic programs. (A) CTP annotation via Over-Representation Analysis (ORA). The heatmap displays the enrichment significance (-log_10_ FDR) of CTP marker genes against MSigDB Hallmark, Gene Ontology (Biological Processes), and Positional gene sets. (B) UMAP embeddings depicting CTP utilization across three independent ccRCC scRNAseq datasets^35–37^. The leftmost column displays harmonized cell-type annotations (B-cell, Benign Epithelium [Benign-Epi], Endothelial, Myeloid and Antigen Presenting Cells [Myeloid/APC], RCC, Stromal, and T and Natural Killer cells [T/NK]; see **Table S3**). Remaining columns display utilization status for select CTPs; utilization is defined as a CTPscore significantly exceeding a gene-wise permuted null distribution (FDR < 0.05; see Methods). Full distribution of CTPs in **Figure S3**. (C) Dot plot displaying the log odds ratio of CTP utilization by cell type, calculated using Fisher’s exact test. Dot size reflects statistical significance after Benjamini-Hochberg correction; dot color indicates the direction and magnitude of enrichment. (D) Schematic of computational processing for RCC tumorgraft RNAseq data^38^, illustrating the separation of human-derived (malignant) and murine-derived (non-malignant) transcripts. (E) Heatmap of CTPscores across 131 RCC tumorgraft models. The bar plot (right) indicates the fraction of tumorgraft lines utilizing each program. (F) Box plots comparing CTPscores in ccRCC versus non-clear cell RCC (nccRCC) tumorgraft lines, with significance assessed by Kruskal-Wallis test. Red point color depicts statistically significant CTP utilization. Abbreviations: FHdRCC, Fumarate Hydratase deficient RCC; pRCC, papillary RCC; tRCC, translocation RCC; uRCC, unclassified RCC.

Because bulk RNAseq of tumor specimens reflects a mixture of cell types, including tumor cells and their TME, we reasoned that individual CTPs could capture tumor-cell-intrinsic (from this point referred to as RCC-intrinsic), tumor-cell-extrinsic processes (i.e., immune or stromal programs), or mixed processes active across multiple cell types (e.g., RNA translation). To test this, we examined CTP utilization in three independent scRNAseq datasets of treatment-naïve ccRCC tumors and matched adjacent normal kidney^35–37^. We harmonized previously defined cell annotations into seven major categories: (1) RCC, (2) B cell, (3) T cell/Natural Killer cells (T/NK cells), (4) fibroblasts and pericytes (Stromal), (5) Endothelial, (6) myeloid and antigen presenting cells (Myeloid/APC), and (7) benign epithelium (Benign-Epi; **Table S3**). We then assigned cell level CTPscores of the programs defined in bulk-RNAseq using the same methodology detailed above. Integrating these results with the over-representation analysis findings allowed us to name each CTP according to two complementary lines of evidence: the cell-type compartment in which the program was predominantly utilized, and, where informative, the dominant biological theme identified by over-representation analysis.

Six CTPs were enriched in RCC cells across all datasets and were designated RCC programs 1–6 (R1–R6). Four CTPs were enriched among TME cell types and were named accordingly: TME-Tcell, TME-Myeloid (TME-Myelo), TME-Endothelial (TME-Endo), and TME-Stroma (**Figure 2B-C, Figure S3, Figure S4**). Two programs were enriched in benign epithelial compartments: B-RenEpi, associated with normal renal epithelium, and B-Epi, which captured broader epithelial signatures, including markers such as *KRT7* (encoding Keratin 7) and *SFTPD* (encoding Surfactant Protein D), potentially reflecting benign non-renal epithelial contamination in metastatic lesions.

Several transcriptomic programs exhibited broad activity across both malignant and non-malignant cells and were termed mixed programs (MP); individual MPs were named according to the biological themes identified by over-representation analysis (**Figure 2A**). The MP-Proliferative (MP-Prolif) demonstrated significant overlap with cell cycle gene-sets and contained proliferation genes such as *CENPA* and *AURKB*. MP-Ychr was associated with Y-chr positional gene sets, composed of genes on the Y-chr (*ZFY, DDX3Y)* and associated with male sex in bulk and scRNAseq datasets (**Figure S5**). Other MPs included MP-Cilia, MP-Translation (MP-Trans), and an uncharacterized program (MP-Unch) which contained a high proportion of non-coding RNAs.

We next examined these programs across a series of 131 RCC tumorgraft lines with available RNAseq data^38^. In tumorgrafts, human non-malignant components such as stroma and innate immune cells are ultimately replaced by murine counterparts^39^.

Through differential alignment of human and murine transcripts to their respective reference genomes, this system enables the isolation of tumor-specific (human) gene expression (**Figure 2D**). Applying CTPscores to RCC tumorgrafts revealed that MP-Prolif was the most frequently utilized program in over 90% of lines. RCC-enriched programs R1–R6 were variably expressed among models and demonstrated histological subtype specificity, with R1–R4 predominantly active in ccRCC and R5–R6 enriched in non-clear cell (nccRCC) variants (**Figure 2E, F**). TME- and benign-epithelial-associated programs were not significantly utilized by tumorgraft lines, suggesting limited RCC-intrinsic expression of these programs.

### RCC consensus transcriptomic programs are associated with canonical driver alterations

ccRCC is characterized by recurring mutations in tumor suppressor genes including *VHL, BAP1, PBRM1, SETD2, PTEN, TSC1,* and others^12^. We sought to determine if CTPs were associated with specific mutations. To this end, we independently tested associations between CTPscores and canonical RCC driver alterations in each dataset. In total, 1,772 samples of the 2,163 in our discovery cohort had matched RNAseq and DNA sequencing data (**Table S1**); mutation frequencies in each dataset are shown in **Figure S6**.

Among RCC intrinsic programs (R1-R6), R1-R4 were positively associated with *VHL* mutations, whereas R5 and R6 were inversely associated with *VHL*. R5 was strongly enriched for *TFE3/TFEB* fusions, and R6 was enriched for *NF2* mutations. The latter association was only significant in the IM151 dataset, perhaps reflecting the low frequency of *NF2* mutations in JR101 (2%) and TCGA (1%) relative to IM151 (6%) (**Figure 3A**). Among the *VHL*-associated programs, R2 was associated with *PBRM1* and *KDM5C* mutations but was inversely associated with *BAP1*. The opposite pattern was observed with R4, which was associated with *BAP1* but inversely associated with *PBRM1*, mirroring the established mutual exclusivity of mutations in these genes^15^. R3 was associated with *PTEN* mutations across all datasets, and *TSC1* mutations in the JR101 and IM151 datasets. The frequency of *TSC1* mutations was 1% in the TCGA dataset versus 6% in JR101 and 10% in IM151, potentially limiting power.

**Figure 3.**
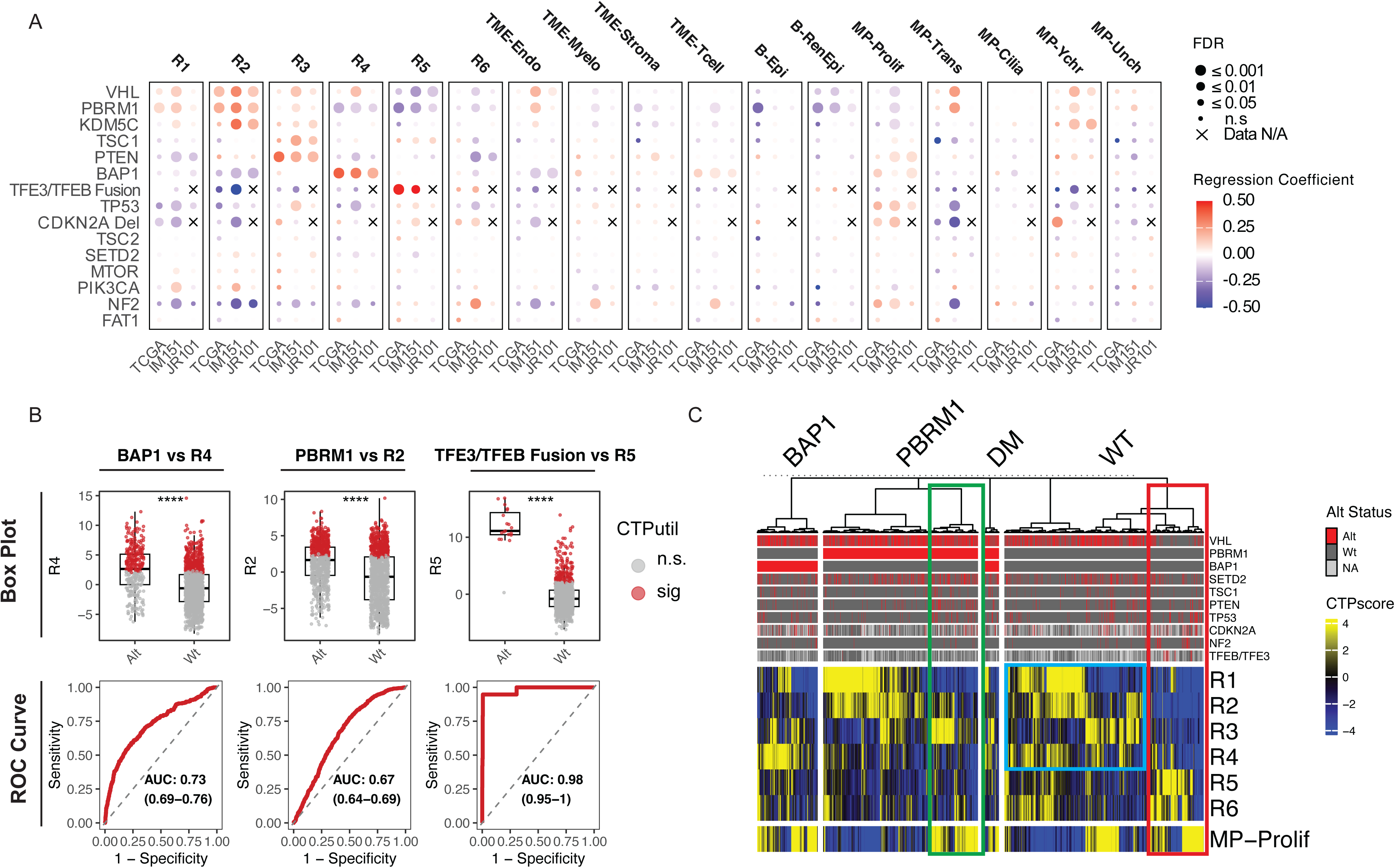
RCC-intrinsic consensus transcriptomic programs are associated with canonical driver alterations. (A) Dot plot displaying the association between CTPscores and canonical driver alterations across the IM151, JR101, and TCGA datasets (n = 1,772 specimens with matched DNA-seq and RNAseq data; see **Table S1**, **Figure S6**). Associations were assessed via logistic regression within each dataset independently. Dot size represents statistical significance (−log₁₀ FDR), and dot color represents the regression coefficient (red, positive association; blue, negative association). An “X” indicates that data were unavailable for a given gene/dataset combination. (B) Box plots (top) of CTPscores stratified by mutation status and Receiver Operating Characteristic (ROC) curves (bottom) evaluating the classification performance of R4 for *BAP1* status, R2 for *PBRM1* status, and R5 for *TFE3/TFEB* fusions. Area under the curve (AUC) and 95% confidence intervals are shown. (C) Heatmap of CTPscores organized by *BAP1/PBRM1* mutation classification: *BAP1*-mutant, *PBRM1*-mutant, *BAP1/PBRM1* double mutant (DM), and wild-type (WT). The top annotation bars indicate the alteration status of key RCC driver genes; samples are arranged by unsupervised hierarchical clustering within each group. Colored boxes depict subclusters of interest; green box, *PBRM1* mutant subcluster enriched in R3 and MP-Prolif; blue box, WT tumors enriched for R1, R2, R3, or R4; red box, WT tumors enriched for *TFE3/TFEB* fusions and *NF2* alterations.

Among mixed programs, MP-Prolif was associated with *BAP1* and *TP53* mutations across all three discovery datasets and with *CDKN2A* deletion where data were available (IM151, TCGA). MP-Ychr was associated with *VHL* and *KDM5C* in the IM151 and JR101 datasets, with the latter relationship potentially reflecting sex-linked effects given that *KDM5C* resides on the X-chromosome and is more frequently mutated in men^40^. Among TME enriched programs, TME-Tcell was associated with *BAP1* mutations in all three datasets, consistent with the “inflamed” microenvironment described for this genotype^19,39^. The remaining TME programs showed inconsistent associations.

We tested these relationships in two validation datasets, the TRACERx Renal study^41^ (TRACERx) and CheckMate 025/010/009^17^, which had matched mutation and RNAseq data available for 230 and 217 specimens, respectively (**Table S4**). Despite a smaller sample size, several associations remained highly significant, including *VHL* with R1-R4, R2 with *PBRM1*, R4 with *BAP1*, and MP-Prolif with *BAP1* and *CDKN2A* loss (**Figure S7**). The association of *PTEN* and *TSC1* with R3 was not statistically significant in this cohort.

In discrimination analyses performed on the combined IM151, TCGA, and JR101 datasets, R2 and R4 distinguished *PBRM1* and *BAP1* mutant versus wild-type tumors with AUCs of 0.67 (95% CI: 0.64-0.69) and 0.73 (95% CI: 0.69-0.76), respectively, whereas R5 identified *TFE3/TFEB* fusion tumors with an AUC of 0.98 (95% CI: 0.95-1) **(Figure 3B)**. The lower AUCs for R2 and R4 compared with the near-perfect performance of R5 indicated that *PBRM1* and *BAP1* mutation status alone did not fully determine program utilization. This could reflect smaller effect sizes, greater within-genotype heterogeneity, or both. To distinguish these possibilities, we examined CTP enrichment within established *BAP1/PBRM1* genotype groups, in which samples are binned into *BAP1* mutant, *PBRM1* mutant, *BAP1/PBRM1* double mutant (DM), and *BAP1/PBRM1* wild type (WT) groups^42,43^.

Stratification revealed that, when comparing *BAP1* and *PBRM1* mutant tumors, R2 and R4 were largely specific to their respective mutation: 44% of *PBRM1* mutant cases utilized R2 versus 5% of *BAP1* mutant cases (p < 0.001), and 59% of *BAP1* mutant cases utilized R4 versus 11% of *PBRM1* mutant cases (p < 0.001). Reciprocally, *PBRM1* mutant cases exhibited higher R1 utilization (45% vs. 21%, p < 0.001), while *BAP1* mutant cases exhibited markedly higher MP-Prolif utilization (56% vs. 27%, p < 0.001, **Figure S7B)**. However, despite these group-level associations, heterogeneity among program utilization was evident among these subgroups. For example, at a high level, *PBRM1* mutant tumors could be stratified into a subset which co-utilized R1 and R2, and another which co-utilized R3 and MP-Prolif (**Figure 3C**, green box). A similar split was observed in *BAP1* mutant cases with most exhibiting dominant MP-Prolif enrichment. There was no significant difference in R3 utilization by mutation subtype.

Among WT tumors, 64% had expression of either R1-R4, which was comparable to other *BAP1/PBRM1* genotype groups, suggesting that a fraction of *BAP1/PBRM1* WT cases may exhibit gene expression resembling mutant cases (**Figure S7C** and **Figure 3C**, blue box). R5 and R6 were highest in WT tumors (**Figure S7B**), with this subset clustering independently within WT tumors (**Figure 3C**, red box). This group was enriched for *TFE3/TFEB* fusions and *NF2* alterations and harbored comparatively fewer *VHL* mutations (**Figure 3C**, red box).

Notably, *TFE3/TFEB* fusion RCC is a distinct histological entity classified separately from ccRCC^8^. Combined with the preferential utilization of R5 and R6 in nccRCC tumorgraft lines and the increased frequency of *NF2* mutations in non-clear cell subtypes^44,45^, these findings suggest that R1-R4 represent transcriptomic states specific to *VHL*-mutant ccRCC, whereas R5 and R6 may capture programs associated with non-ccRCC histologies.

### Consensus transcriptomic programs reveal BAP1- and PBRM1-associated trajectories which converge on a shared aggressive phenotype

ccRCC driver mutations accumulate in a sequential and branching pattern, with biallelic *VHL* inactivation as the initiating event followed by divergence into either *PBRM1*- or *BAP1*-mutant lineages^9–12^. Whereas *BAP1-*mutant ccRCC tend to be more aggressive than *PBRM1-*mutant counterparts, *PBRM1-*mutant ccRCC frequently acquire additional mutations resulting in more aggressive biology. These include mutations in mTOR pathway genes (including *PTEN* and *TSC1),* and deletions in chromosome 9p (containing *CDKN2A),* among others^10,11^. Given the heterogeneous utilization of R1-R4 and MP-Prolif in *BAP1/PBRM1* mutant tumors, and the association of these CTPs with the canonical drivers above, we asked whether relationships among these programs could be organized into a coherent transcriptomic trajectory mirroring this genomic progression.

Trajectory inference methods, such as diffusion mapping, have been widely applied to scRNAseq data to order cells along continuous progression axes, but their application to bulk RNAseq is limited by the confounding influence of tumor microenvironment composition. Because our CTP framework deconvolves RCC-intrinsic from tumor-cell-extrinsic signals, we reasoned that diffusion mapping applied to RCC-intrinsic CTPscores, rather than raw bulk expression, would isolate variation in tumor cell transcriptional states and enable trajectory inference across large bulk RNAseq cohorts. To test this, we applied destiny^46^, a manifold learning approach that embeds samples in a low-dimensional space based on transition probabilities, enabling inference of continuous progression axes (**Methods**). The analysis was applied to 1,463 (70.0%) of 2,091 tumor specimens that significantly utilized at least one *VHL*-associated program (R1–R4), using CTPscores for the five RCC-intrinsic programs associated with canonical drivers (R1, R2, R3, R4, and MP-Prolif) as input. Note, MP-Prolif was retained despite its mixed designation, given its high expression evidence in RCC tumorgrafts (suggesting RCC-intrinsic expression) and its association with *CDKN2A* and *TP53*, genomic drivers that occur late in RCC evolution. The remaining 628 specimens were most commonly dominated by TME programs (35%) or MP-Prolif (27%), with smaller subsets dominated by R5/R6 (12%), benign epithelial programs (10%), other mixed programs (11%), or no significantly utilized CTP (5%; **Figure S8**). These cases were excluded from trajectory inference because their RCC-intrinsic state could not be unambiguously placed along the *VHL-*associated axis.

The first axis of the resulting embedding (diffusion component 1, DC1) defined a continuum that placed R1 at the origin, R2 and R4 in the center, and R3 and MP-Prolif at the terminus (**Figure 4A**). To orient this continuum and test whether it reflected biological progression, we asked whether DC1 was associated with histopathological aggressiveness. Indeed, the scaled DC1 coordinate (hereby referred to as pseudotime) was associated with increasing Fuhrman nuclear grade, a histopathological measure of tumor aggressiveness, supporting the directionality of the inferred trajectory (**Figure 4B**). Consistent with this orientation, several R3 marker genes have previously been associated with aggressive features in ccRCC, including *CP* (ceruloplasmin), *PTHLH* (encoding PTHrP), and *IL6* (**Figure 4C**)^47–50^.

**Figure 4.**
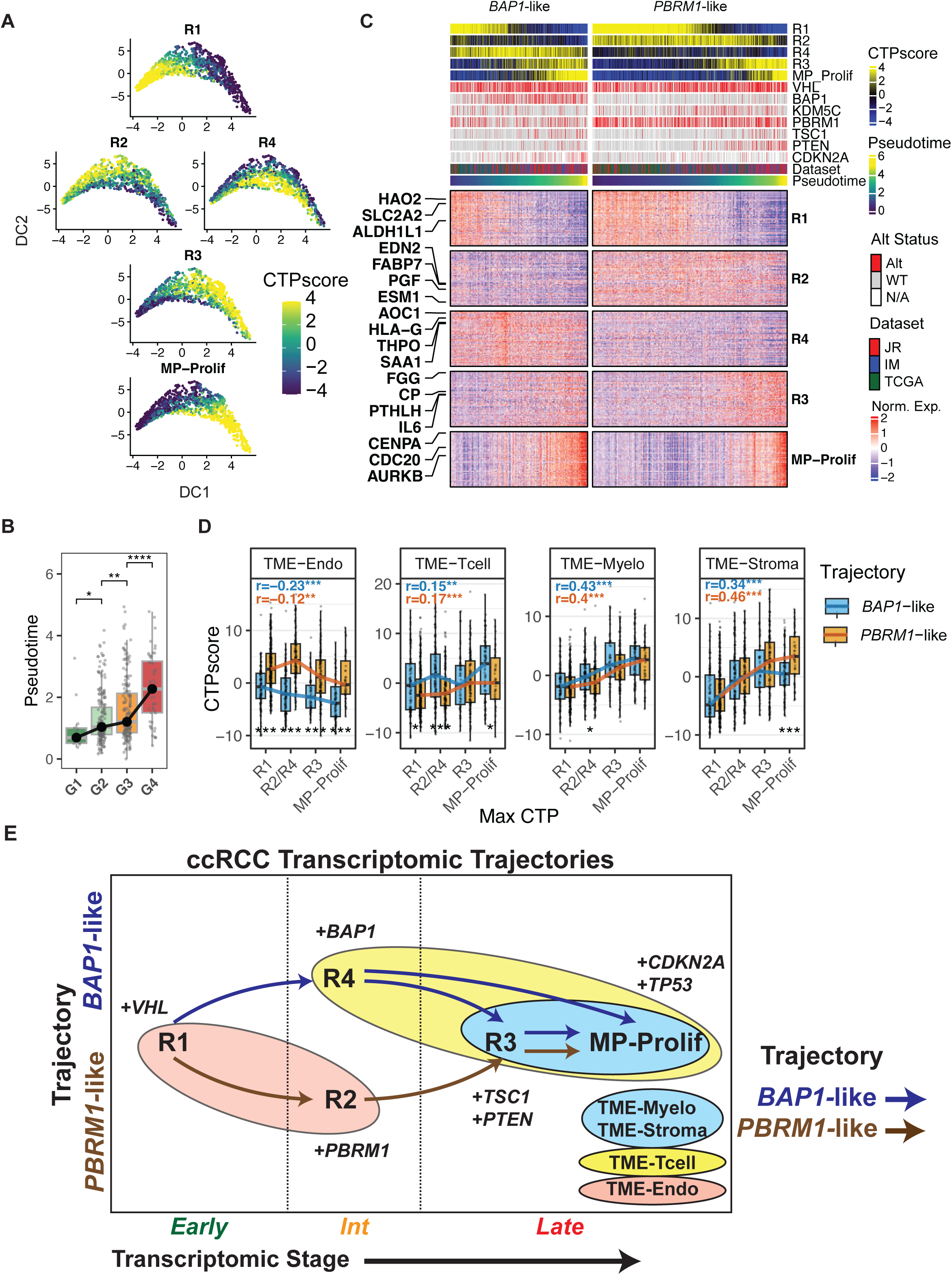
*BAP1*- and *PBRM1*-associated transcriptomic trajectories converge on a shared aggressive phenotype. (A) Diffusion map depicting putative transitions between ccRCC-intrinsic programs (R1–R4 and MP-Prolif). Analysis was performed using the destiny algorithm on 1,463 (70%) bulk RNAseq specimens utilizing at least one of R1-R4. Faceted scatter plots display the first two diffusion components (DC1 vs. DC2), with individual points representing tumor specimens colored by CTPscore. (B) Heatmap of expression for CTP marker genes and corresponding CTPscores, with samples ordered by pseudotime and split into two inferred trajectories defined as *BAP1*-like (R4 > R2) and *PBRM1*-like (R2 > R4). Top annotation bars indicate driver alteration status, dataset of origin, pseudotime, and CTPscores. (C) Box plots of pseudotime stratified by Fuhrman nuclear grade (G1–G4) in TCGA (n = 522). p values reflect Wilcoxon rank-sum tests. (D) Tumor microenvironment (TME) program scores (TME-Endo, TME-Tcell, TME-Myelo, TME-Stroma) stratified by trajectory (*BAP1*-like vs. *PBRM1*-like) and dominant RCC-intrinsic program (R1, R2/R4, R3, MP-Prolif). Trend statistics (brown and blue line) reflect Spearman correlation between ordered CTP max state (R1 → R2/R4 → R3 → MP-Prolif) and TME score within each trajectory. Spearman rho and p value reported for each trend. Pairwise comparisons of trajectories at each CTP state by Wilcoxon rank-sum tests. (E) Schematic summarizing the two transcriptomic trajectories (*BAP1*-like and *PBRM1*-like) and transcriptomic stage (TS), depicting progression from an Early state (R1) through intermediate branching (R2 vs. R4) to a Late convergent state (R3/MP-Prolif), annotated with associated driver alterations and TME features.

We further examined the placement of R1 at the pseudotime origin using scRNAseq, where we noted that R1 was utilized in both ccRCC cells and benign renal epithelium **(Figure 2C)**. Detailed inspection revealed a high fraction of R1 marker genes were expressed in both proximal tubular cells and ccRCC cells, but not in non-proximal tubular benign renal cells. In contrast, the B-RenEpi program sharply distinguished benign renal epithelium from tumor, with high expression in benign renal epithelial cells (both proximal tubular and non-proximal tubular cells) and minimal expression in R1-utilizing ccRCC cells (**Figure S9**). These observations suggest that R1 captures proximal-tubule-specific genes which are retained at initial malignant transformation but ultimately down-regulated with tumor progression towards more aggressive disease, supporting placement of R1 at the pseudotime origin.

Whereas DC1 defined the primary progression axis, DC2 and DC3 resolved a branch point in which tumors transitioned from an R1-enriched early state toward either an R2-enriched or R4-enriched intermediate state before converging on R3- and MP-Prolif-enriched late states (**Figure 4A**; **Figure S10**). This indicated two converging trajectories defined by their dominant intermediate state: *PBRM1*-like (R2 CTPscore > R4 CTPscore) and *BAP1*-like (R4 CTPscore > R2 CTPscore). The branching and converging structure is further supported by co-utilization analysis (**Figure S10B**). R1 was co-utilized with both R2 and R4, but R2 and R4 were mutually exclusive (p < 0.001 for both comparisons, Fisher’s exact test). Likewise, R3 co-occurred with R2 and R4, but was mutually exclusive with R1, and MP-Prolif co-occurred with R3 (p < 0.001 for all comparisons). Ordering tumors by pseudotime and stratifying by trajectory revealed smooth transitions in CTPscore and gene expression along each trajectory (**Figure 4C**).

To formalize and test this framework, we organized these observations into two complementary concepts: *trajectory*, defined by the relative magnitude of R2 versus R4 CTPscores (*PBRM1*-like when R2 > R4, *BAP1*-like when R4 > R2), and *transcriptomic stage* (TS), defined as the position along the progression axis. TS stratifies tumors into three levels; Early (R1-dominant), Intermediate (R2- or R4-dominant), and Late (R3- or MP-Prolif-dominant), with each sample assigned a TS based on its maximally enriched CTP. Conceptually, trajectory and TS describe distinct aspects of tumor progression: trajectory specifies which path a tumor follows, while TS specifies how far along that path the tumor has progressed. By analogy, the two trajectories can be thought of as distinct routes a tumor may take, while TS indicates how far along that route the tumor has traveled.

We next asked how TME composition varied along this two-axis progression model: both along TS (early to late) and between trajectories (*BAP1*-like vs. *PBRM1*-like). We compared TME-Endo, TME-Tcell, TME-Myelo, and TME-Stroma CTPscores across samples organized by their TS and trajectory (**Figure 4D**). The most pronounced difference between trajectories was observed in the TME-Tcell and TME-Endo programs. The *PBRM1*-like trajectory exhibited higher TME-Endo scores across all TS categories (p < 0.001), whereas the *BAP1*-like trajectory exhibited greater enrichment for TME-Tcell programs, with trajectory differences most evident at the Intermediate stage (R2 vs. R4). This pattern may reflect the distinct composition of the intermediate programs. R2 contains pro-angiogenic genes including *EDN2*, *PGF*, and *ESM1*, whereas R4 contains non-classical immune checkpoint molecules (*HLA-G*, *HHLA2*), acute phase markers (*SAA1*, *SAA2*), and interferon-stimulated genes (*GBP7*), suggesting a distinct immune-modulated state (**Table S2**). Across both trajectories, TME-Myelo and TME-Stroma scores increased in a stepwise fashion from Early to Late TS (p < 0.001 for all comparisons), consistent with the composition of R3, which encodes mediators of myeloid recruitment and activation (e.g., *IL6*, *CXCL5*) matrix metalloproteinases (e.g., *MMP1*, *MMP12)*, and immunosuppressive metabolic enzymes (e.g., *ARG2, KMO*) (**Table S2**).

Together, these findings suggest a model in which ccRCC tumors progress along one of two transcriptomic trajectories, *PBRM1*-like or *BAP1*-like, that diverge at an intermediate state and converge on a shared aggressive Late state (**Figure 4E)**. The *PBRM1*-like and *BAP1*-like trajectories are distinguished by their Intermediate states (R2- and R4-enriched, respectively) and by trajectory-specific TME features (vascular for *PBRM1*-like, T cell-enriched for *BAP1*-like), whereas the convergent Late state is characterized by R3 and MP-Prolif utilization, increased nuclear grade, and progressive myeloid and stromal infiltration along both trajectories. This framework links the branching genomic architecture of ccRCC to a corresponding branching transcriptomic architecture and proposes TS and trajectory as orthogonal axes for describing tumor progression.

### Spatial transcriptomics and multiregional sequencing enable detection of intra-tumoral transcriptomic stage transitions

We next tested whether the branching trajectories inferred from bulk tumors could be detected intra-tumorally using complementary sampling strategies that resolve intratumoral heterogeneity. We first analyzed Visium spatial transcriptomic data from eight nephrectomy specimens with adjacent conventional clear cell (CC) and sarcomatoid-dedifferentiated (Sarc) regions, deposited by Salgia et al.^51^ Because sarcomatoid dedifferentiation represents an aggressive, late state of ccRCC, this setting provides an internal reference for directional change. We assigned spot-level TS and trajectory using the same definitions applied to bulk tumors and compared their distributions by histologic annotation. This revealed a significant shift from predominantly Intermediate TS in CC regions to predominantly Late TS in Sarc regions (**Figure 5A**; p < 0.001). The trend persisted after stratifying by subject; most CC regions contained a higher fraction of Early or Intermediate TS relative to matched Sarc regions. Although the effect was modest in three cases, in no case did Sarc regions contain a higher fraction of Early TS spots than the paired CC region (**Figure S11**).

**Figure 5.**
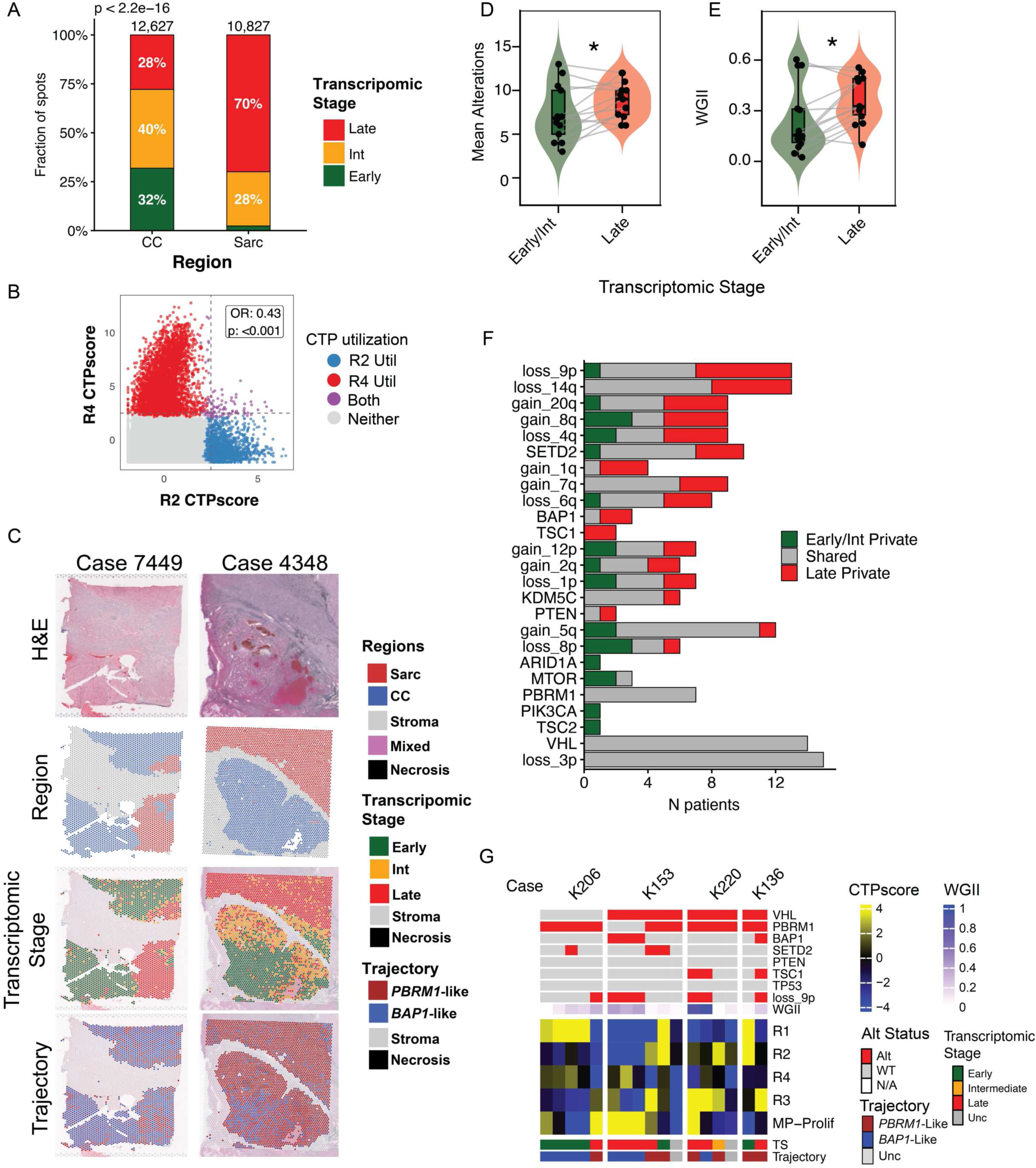
Spatial and multi-regional profiling capture intra-tumoral progression along transcriptomic stage. (A) Stacked bar plot of Visium spot counts across clear cell (CC) and sarcomatoid (Sarc) regions from eight nephrectomy specimens^51^, stratified by transcriptomic stage (TS: Early, Intermediate, Late). p value reflects chi-squared test. (B) Scatter plot of R2 versus R4 CTPscores at the spot level, colored by utilization status (R2 only, R4 only, both, or neither). Odds ratio and p value from Fisher’s exact test are shown. (C) Representative spatial mapping of two cases (Case 7449 and Case 4348) displaying, from top to bottom: H&E histology, tissue regions (CC, Sarc, Stroma, Mixed, Necrosis), spot-level TS assignment (Early, Intermediate, Late), and trajectory (*PBRM1*-like vs. *BAP1*-like). (D-E) Violin and overlaid box plot comparing the mean number of somatic alterations and whole-genome instability index (WGII) between Early/Intermediate and Late regions in the TRACERx cohort. p value reflects a Wilcoxon rank-sum test. (F) Per-alteration partitioning of driver mutations and SCNAs across the 15 TS-Discordant TRACERx Renal patients (those with both Early/Intermediate-TS and Late-TS regions). Each alteration is classified per patient as Early/Int Private (green), Shared (gray), or Late Private (red), based on its regional distribution. Alterations are ordered by Late-Private frequency; events absent across all 15 patients are omitted. (G) Multiregional profiling of four representative TRACERx patients (K206, K153, K220, K136). For each patient, dendrograms from unsupervised hierarchical clustering (top) are displayed alongside heatmaps of driver alteration status, WGII, TS, trajectory, and CTPscores for R1–R4 and MP-Prolif across sampled regions.

We next assessed trajectory and found that three cases exhibited a predominantly *PBRM1*-like trajectory (R2-high/R4-low), three exhibited a *BAP1*-like trajectory (R4-high/R2-low), and two showed a mixture across spots **(Figure S11).** In Case 4352, these trajectories were spatially segregated, with the *PBRM1*-like trajectory restricted to the Sarc region and the *BAP1*-like trajectory enriched in the CC region. In Case 7447, R2 and R4 were intermixed across both regions, potentially reflecting reduced discriminatory power of R2/R4 when both programs have low enrichment (**Figure S11**). Nevertheless, at the spot level, R2 and R4 utilization were strongly mutually exclusive (**Figure 5B**).

Spot-level co-utilization patterns were largely concordant with patterns in bulk data and further supported the inferred trajectory ordering. R1 co-occurred with both R2 and R4, whereas R2 and R4 were mutually exclusive, consistent with a branching intermediate state. R1 and R3 were also mutually exclusive, supporting an intermediate step between these programs (p < 0.001 for all comparisons; **Figure S11**). One pattern, however, differed between the two data modalities. In bulk data, R3 co-occurred with both R2 and R4, and MP-Prolif co-occurred with R3 (**Figure S10**). At the spot level, R3 was instead mutually exclusive with R4, while MP-Prolif co-occurred with R4 and was mutually exclusive with R2. This raises the possibility of a direct transition from R4 to MP-Prolif that bypasses R3 in the *BAP1*-like trajectory; whether an analogous *PBRM1*-like transition from R2 to MP-Prolif exists could not be determined from the present data, given the limited number of cases.

The spatial relationship among trajectory, TS, and sarcomatoid dedifferentiation is highlighted by Cases 7449 and 4348 (**Figure 5C**). In these cases, CC regions contained a mixture of Early and Intermediate TS, whereas Sarc regions were predominantly Late. In contrast, trajectory assignments were largely homogeneous, with *BAP1*-like trajectory in Case 4348 and *PBRM1*-like trajectory in Case 7449. The spatial distribution of TS, trajectory, and CTPscores is shown for all samples in **Figures S11 and S12**.

To connect trajectories and TS to driver mutation accumulation within patients, we next assigned trajectory and TS to the TRACERx cohort, which includes multiregional RNA and DNA sequencing from at least two specimens for 64 patients (n = 213 specimens)^41^. Patients were grouped by the TS assignments of their sampled regions: TS Early/Intermediate (all regions Early or Intermediate), TS Late (all regions Late), and TS Discordant (Early/Intermediate to Late transitions across regions, n = 15). Driver mutations, CTPscores, TS, and trajectory assignments for all patients are shown in **Figure S13A-C**. We next examined how frequently trajectory assignments were shared across regions within a patient. Trajectory was concordant in 63% of cases, significantly higher than a bootstrapped null in which trajectories were randomly assigned (p < 0.001; **Figure S13D**). Whole-genome instability index (WGII) and the number of driver alterations increased stepwise across TS, and this relationship persisted in paired within-patient comparisons among 15 patients with discordant TS (Early/Intermediate versus Late regions; **Figure 5D–E, Figure S13E-F**).

Among these 15 patients with discordant TS, the most common private alterations acquired in Late regions were loss of 9p (containing *CDKN2A*) and loss of 14q in 6/15 and 5/15 patients, respectively (**Figure 5F**). For example, shifts from Early to Late TS aligned with 9p loss in K206 and with acquisition of *TSC1* alterations plus 9p deletion in K220 and K136. The relationship among mutations, TS, and trajectory was particularly well illustrated by K153: three regions harboring *VHL/BAP1* mutations with 9p loss were Late TS with a *BAP1*-like trajectory, whereas distinct regions from the same tumor with *VHL/PBRM1* mutations exhibited earlier TS along the *PBRM1*-like trajectory (**Figure 5G**).

We next extended this comparison to the cohort level by relating TS to the genomic evolutionary subtypes (EvoTypes) previously defined for the TRACERx Renal cohort^11,41,52^, which classify tumors by their constellation of truncal and subclonal alterations. The *VHL*-mono EvoType, characterized by isolated *VHL* inactivation and a more indolent clinical course, contained the highest fraction of Early TS regions (4/6 patients, 67%). In contrast, the *BAP1*-driven and multiple-clonal-driver EvoTypes (the latter reflecting tumors with multiple truncal driver mutations) were enriched for Late TS (4/8 patients, 50% and 8/12 patients, 75%, respectively), consistent with their more aggressive clinical behavior (**Figure S13G**).

### Transcriptomic stage is associated with clinical outcomes in RCC

We next evaluated whether TS, inferred trajectory, and individual CTP utilization were associated with clinical outcomes across three settings: TCGA (predominantly localized disease; overall survival) and two phase III frontline metastatic trials, IM151 and JR101 (progression-free survival; **Figure 6**, **Table S1**).

**Figure 6.**
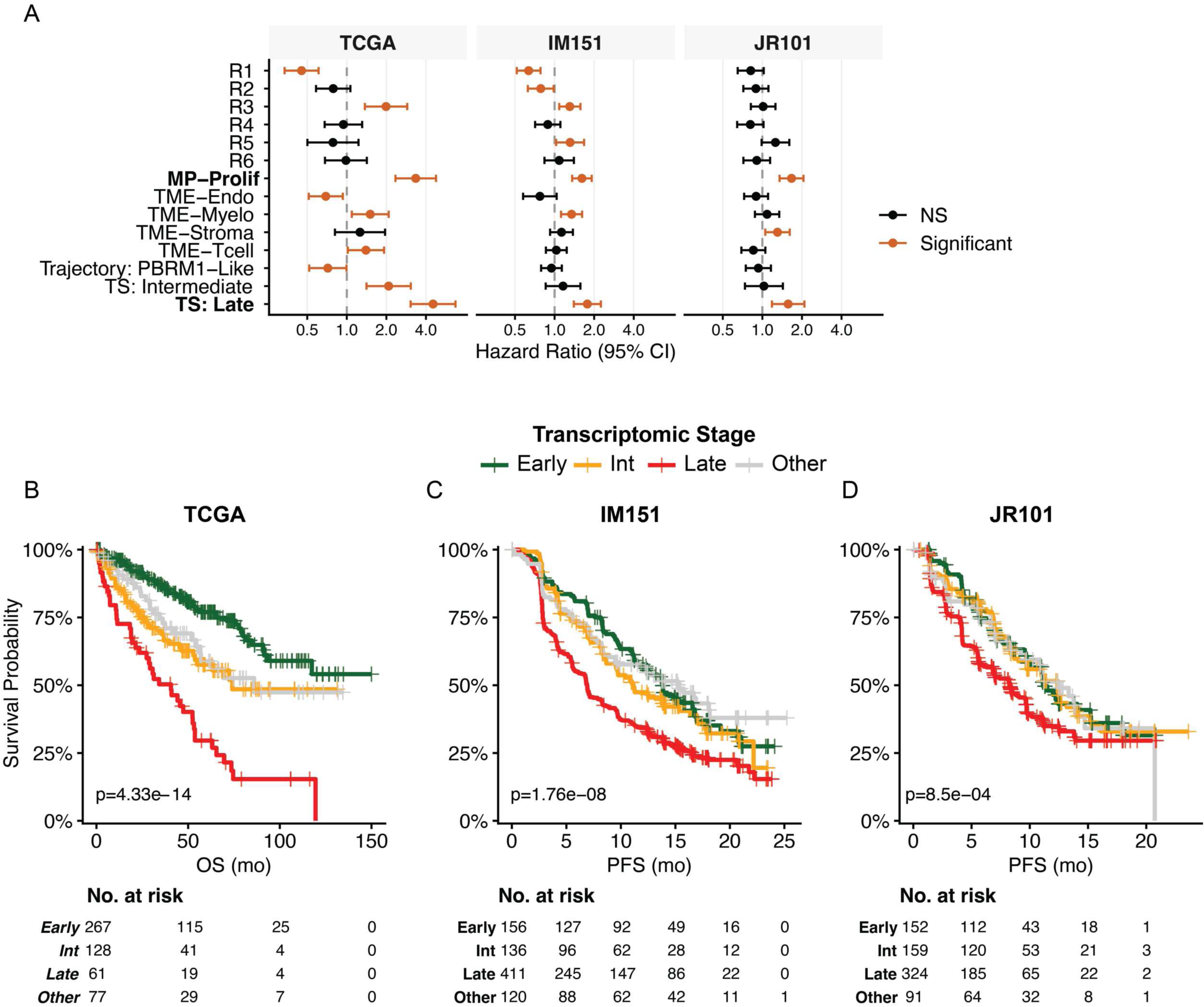
Transcriptomic stage is prognostic in ccRCC. (A) Forest plots of univariate hazard ratios (HRs) for individual CTP utilization, trajectory assignment, and transcriptomic stage (TS) across three independent cohorts: TCGA (overall survival [OS]; n = 542), IM151 (progression-free survival [PFS]; n = 823), and JR101 (PFS; n = 726). HRs represent risk of death (TCGA) or risk of progression or death (IM151, JR101); points denote HRs and horizontal bars denote 95% confidence intervals. Significant associations are highlighted in orange. (B–D) Kaplan-Meier survival curves stratified by TS (Early, Intermediate, Late; “Other” shown in gray) for OS in TCGA (B) and PFS in IM151 (C) and JR101 (D). Log-rank p values and numbers at risk are displayed below each plot.

The distribution of TS differed substantially between TCGA and the metastatic trial datasets, with TCGA containing a larger proportion of Early tumors. This largely reflected the high fraction of stage I-II disease in TCGA, which was enriched for Early TS. In contrast, in IM151 and JR101, all patients had metastatic disease at the time of sampling or subsequently developed metastatic disease (**Figure S14A-B**).

Across all three datasets, Late TS was consistently associated with inferior outcomes in univariate analyses, with the largest effect sizes observed in TCGA, potentially reflecting longer follow-up. In multivariable models, TS remained associated with outcome after adjustment for established clinical covariates (stage and grade in TCGA; treatment arm and IMDC risk where available in IM151) supporting TS as an independent prognostic correlate (**Figure 6, Tables S5-S7**). Several TME-associated programs were associated with outcome in TCGA (e.g., TME-Tcell and TME-Myelo with poorer outcomes; TME-Endo with more favorable outcomes), but these associations were attenuated in the metastatic trial cohorts.

The *BAP1*/*PBRM1* genomic classifier has been previously associated with outcomes in the TCGA and other cohorts with predominantly localized disease^13,42,43^. We therefore asked whether TS would remain prognostic in a model additionally controlling for *BAP1*/*PBRM1* mutation status. In the TCGA, TS remained highly prognostic after adjustment for stage, grade, and *BAP1*/*PBRM1* classifier, with a HR of 3.47 (95% CI 1.87–6.43) for Late vs. Early and 2.27 (95% CI 1.24–4.15) for Intermediate vs. Early (**Table S8**). Notably, in this model the only other covariate reaching statistical significance was stage IV disease; neither nuclear grade nor *BAP1*/*PBRM1* status was independently significant.

We next examined the distribution of TS and trajectory by *BAP1/PBRM1* genotype. Consistent with our trajectory definitions, *BAP1*-mutant tumors were strongly enriched for the *BAP1*-like trajectory (83%) and *PBRM1*-mutant tumors for the *PBRM1*-like trajectory (67%), supporting the genotype-trajectory association (**Figure S14C**). *BAP1*-mutant tumors were enriched for Late and Intermediate TS relative to *PBRM1*-mutant tumors (49% Late and 37% Intermediate in *BAP1* versus 35% late and 16% Intermediate in *PBRM1*), in keeping with their more aggressive clinical behavior^42,43^ (**Figure S14D**). However, the TS distribution within PBRM1-mutant tumors was strongly dataset-dependent: *PBRM1*-mutant tumors were predominantly Early TS in TCGA (68%) but predominantly Late TS in IM151 and JR101 (43% and 50%, respectively), mirroring the broader cohort-level shift toward Late TS in the metastatic settings.

Trajectory assignments, on the other hand, were largely stable across datasets (**Figure S14E-F**). This suggests that while trajectory (*BAP1*-like vs. *PBRM1*-like) is largely fixed by genotype, TS varies with clinical disease stage, consistent with TS capturing tumor progression along a genotype-defined path.

We next tested whether TS or specific CTPs interacted with treatment arm (i.e., predicted differential benefit) in IM151 and JR101. IM151 compared atezolizumab plus bevacizumab with sunitinib, whereas JR101 compared axitinib plus avelumab with sunitinib; both were conducted in the treatment-naïve metastatic setting. TS did not show a consistent interaction with treatment arm. In contrast, R1 utilization demonstrated a consistent interaction across both studies, with MP-Prolif nearing significance in IM151 and strongly significant in JR101 (**Figure S15**). Stratified Kaplan–Meier analyses illustrated these effects: patients whose tumors utilized R1 derived limited benefit from immune checkpoint inhibitor therapy combinations relative to sunitinib monotherapy, whereas patients whose tumors utilized MP-Prolif exhibited greater relative benefit from immune checkpoint inhibitor combinations versus sunitinib monotherapy (**Figure S14**).

## Discussion

Here, we present a framework for identifying and contextualizing consensus transcriptomic programs (CTPs) from bulk RNAseq when multiple large cohorts exist for a single disease, and we apply it to ccRCC. We define 17 CTPs across three independent datasets with varying degrees of co-utilization at the sample level; a key distinction from prior transcriptomic subclassification efforts in ccRCC^27,53,54^, which assign tumors to discrete subtypes. We find that CTPs are associated with distinct cellular sources, driver alterations, histopathological grade, and clinical outcomes.

Moreover, we find that a subset of CTPs assemble into converging trajectories that integrate gene expression with genotype, histological features, TME composition, and clinical outcomes.

NMF has long been used to uncover coordinated gene expression structure, but conventional applications typically require selecting a single factorization rank within a single cohort, leaving open questions of rank dependence and overfitting^22^. Moreover, this reliance on single rank disregards the hierarchical organization of biological systems and prior work demonstrating that applying unsupervised learning across multiple ranks can recover distinct biological features at different resolutions of factorization^22,26,55,56^. We addressed these issues by identifying recurrent programs across multiple independent ccRCC datasets, leveraging cross-dataset consensus as a constraint that favors biological signal over artifact. Whereas a prior consensus NMF approach was developed to stabilize the stochastic output of NMF at a given rank by aggregating across random initializations^57^, our framework systematically explores a range of factorization ranks and consolidates recurrent structure, simultaneously reducing dependence on any single rank and increasing sensitivity to processes that emerge at particular resolutions of factorization. We further use a gene-wise permutation-derived null to scale program scores, providing a principled baseline for interpreting whether a program is meaningfully utilized within a sample rather than only relative to other samples in the same cohort. By anchoring usage to a null distribution, we enable both comparisons across datasets and downstream association analyses that treat each program as an independent quantitative trait. A second methodological contribution is the application of pseudotime analysis to bulk RNAseq cohorts. While pseudotime trajectories are widely used for single-cell analysis^46,58^, pseudotime methods applied to bulk gene expression leave the trajectory space susceptible to TME-driven variation ^49^. By restricting the input to RCC-intrinsic CTPs, we ground the inferred trajectory in tumor cell transcriptional states rather than microenvironmental composition. Nevertheless, integrating CTPs with PhenoPath’s covariate-based pseudotime framework represents a natural extension, particularly for large cohorts with heterogeneous clinical or genomic phenotypes where formal covariate modeling could yield additional resolution. We provide an R package, *rC3TP*, that applies CTP weights, computes null-scaled scores, and assigns trajectory and TS in user-provided gene expression data, enabling reproducible scoring of these signatures across datasets.

A central biological finding of this study is that ccRCC driver-associated programs (R1–R4 and MP-Prolif) are organized by trajectory analysis into two branching then converging trajectories that mirror established genomic models of ccRCC evolution.

Cellular attribution of CTPs through scRNAseq and tumorgraft data was essential to this interpretation, allowing RCC-intrinsic programs to be examined separately from microenvironmental signal. Our trajectory framework resolves the RCC-intrinsic landscape along two axes: (i) a transcriptomic stage axis that tracks progression from early, lower-grade, favorable-prognosis states toward late, high-grade, poor-prognosis states, and (ii) a trajectory axis that distinguishes intermediate branching states aligned with *BAP1*-like versus *PBRM1*-like lineages. The early end of TS (R1) captures a proximal-tubule lineage state that is downregulated with progression, while the late end (R3 and MP-Prolif) capture gene expression previously linked to poor outcomes; the intermediate branching mirrors the mutual exclusivity of *BAP1* and *PBRM1* mutations.

Spatial transcriptomic analysis and multiregional sequencing provided convergent intra-tumoral support: Late TS co-localized with sarcomatoid dedifferentiation, and TS progression was associated with increasing driver alteration burden and whole-genome instability, in some patients aligning with the acquisition of private alterations such as 9p deletion or *TSC1* mutations.

An important corollary is that mutation status alone incompletely predicts transcriptional state. We observe discordance in both directions: tumors harboring a driver alteration but lacking the expected transcriptomic program, and tumors without detectable mutations exhibiting a transcriptomic state resembling the corresponding lineage. These observations are consistent with a model in which RCC-intrinsic transcription reflects the integrated consequence of multiple genomic events, epigenetic state, and microenvironmental context, and in which distinct molecular routes can converge on similar expression states. Such phenocopying may arise from undetected or subclonal genomic events, alternative drivers affecting convergent pathways, epigenetic reprogramming, or microenvironmental signaling. These possibilities are not mutually exclusive and represent avenues for future mechanistic dissection.

This framework also provides insight into tumor microenvironment remodeling in ccRCC. Classically, the ccRCC TME has been described as either inflamed or angiogenic^19,39^; our approach recovers angiogenesis-associated (TME-Endo) and T cell-associated (TME-Tcell) programs, alongside myeloid (TME-Myelo) and stromal (TME-Stroma) programs. We found that TME composition was influenced by both trajectory and TS. *PBRM1*-like tumors were relatively enriched for TME-Endo in earlier/intermediate states, *BAP1*-like tumors for TME-Tcell programs, while TME-Myelo and TME-Stroma increased stepwise with TS along both trajectories. This structure offers a parsimonious explanation for discrepant reports linking *PBRM1* mutations to either angiogenic or inflamed microenvironments^16–20^. Cohorts differing in their TS distribution, for example localized cohorts enriched for Early/Intermediate tumors versus metastatic or refractory cohorts enriched for Late tumors, may yield different conclusions when comparing on *PBRM1* mutation status alone. In this view, TS acts as a hidden stratifier that modulates genotype-TME associations and may explain why mutation-based predictors of therapy response have shown context dependent results.

These observations have several translational implications. First, TS provides a quantitative axis that may refine risk stratification in localized disease beyond conventional histopathologic grade and stage. This is particularly relevant in the adjuvant setting, where pembrolizumab is approved following nephrectomy based on improved disease-free and overall survival, yet the absolute benefit is modest and many patients are cured by surgery alone^7^. Patients with Early TS tumors, which exhibit favorable biology, could be spared unnecessary immunotherapy exposure, while patients with Late TS tumors may warrant intensified adjuvant strategies or closer surveillance. Second, in the metastatic setting, R1 and MP-Prolif utilization interacted with treatment arm across two independent phase III trials (IM151 and JR101). Patients whose tumors utilized R1 derived limited additional benefit from ICI/VEGF combination therapy relative to sunitinib alone, whereas patients whose tumors utilized MP-Prolif exhibited greater relative benefit from combination therapy. Unexpectedly, TME-associated programs did not show consistent treatment interactions, suggesting that the predictive signal resides in RCC-intrinsic biology rather than microenvironmental composition alone. The OPTIC trial is prospectively testing a biomarker-directed strategy based on IM151-derived transcriptomic subtypes (NCT05361720)^59^, but those subtypes were not predictive when applied retrospectively to JR101^29^, highlighting a potential limitation of classifiers that conflate multiple biological processes into a single label. By disentangling individual programs contributing to such composite subtypes, our approach identifies specific RCC-intrinsic programs with treatment interactions (R1, MP-Prolif) that may have been obscured within broader subtype definitions.

Several limitations should be acknowledged. Our approach, like other matrix factorization methods, is optimized to capture linear structure and may miss non-linear dependencies. Bulk RNAseq inevitably conflates malignant state with cell-type composition; although single-cell and tumorgraft mapping provide strong support for cellular attribution, residual confounding may influence specific associations. Tumor-extrinsic programs in bulk remain coarse relative to the diversity of immune and stromal subsets resolvable by scRNAseq. Finally, the trajectory and translational implications proposed here are based on associations across cohorts and orthogonal modalities; functional validation through mechanistic experiments will be needed to establish causal relationships between specific CTPs, TS advancement, and treatment response.

Despite these limitations, the convergence of evidence across independent cohorts, sequencing modalities, and intra-tumor datasets supports the robustness of the core model. By defining a set of recurrent ccRCC transcriptomic programs and quantifying their usage, we identify two branching ccRCC transcriptomic trajectories that converge on a shared aggressive late state, providing a bridge between genotype, transcriptomic stage, microenvironmental remodeling, and clinical outcome. More broadly, this work illustrates how consensus program discovery across multiple cohorts can yield robust and interpretable transcriptional axes that are well suited to mechanistic interrogation and translational biomarker development.

## RESOURCE AVAILABILITY

**Lead Contact:** Requests for further information and resources should be directed to and will be fulfilled by the lead contact, Roy Elias.

### Materials Availability

This study did not generate new unique reagents.

### Data and code availability

Processed bulk RNAseq datasets and accompanying mutation and clinical annotations are publicly available per the original studies, including the TCGA^9^, IM151^27^, JR101^5^, UTSW RCC Tumorgraft platform^38^, Checkmate 025/010/009^17^, and TRACERx cohort^41^. Likewise, processed single-cell and bulk RNAseq datasets are publicly available per the original publication; Salgia et al.^51^, Li et al.^35^, Zhang and Narayanan et al.^36^, and Alchahin and Mei et al^37^. All code and source data for every figure are deposited on GitHub (https://github.com/rmelias2/rcc_ctp_manuscript ). To facilitate application of these programs to external datasets, we developed an R package (rC3TP; https://github.com/rmelias2/rC3TP ) that computes sample level CTPscores, CTP utilization, trajectory and transcriptomic stage on user-supplied expression matrices using the workflow defined in this manuscript.

## Acknowledgements

The authors wish to thank Alexander V. Favorov, Loyal Goff, Jeanette Johnson, Shahin Mohammadi, and Thomas Sherman for feedback on multi-dimensional factorization methods and Rachel Karchin, Jiaying Lai, and Laura Wood for integrated mutational and gene expression analysis techniques. Funding was provided by NIH/NCI grants U24CA284156 (G.S-O’B, E.J.F, A.D), U01CA253403 (E.J.F), U01CA271273 (E.J.F.), P30CA006973 (S.Y.), U54CA274370 (S.Y.), P50CA272391 (S.Y., R.E.); Maryland Cancer Moonshot Research Grant to the Johns Hopkins Medical Institutions (FY24) (A.D. and E.J.F.); Break Through Cancer (R.E., E.J.F., and A.D.), the Commonwealth Foundation (S.Y.), and the V Foundation for Cancer Research (S.Y.).

## METHOD DETAILS

### Defining Consensus Transcriptomic Programs

Transcripts per million (TPM) expression matrices were obtained for each of the three discovery cohorts, which differed in tissue handling, library preparation, and alignment pipelines. Full details of RNA extraction, library preparation, and quantification for each cohort are described in the respective primary publications and GDC documentation^5,27,62^. Briefly, for the TCGA clear cell renal cell carcinoma (KIRC) dataset, RNAseq was performed on fresh-frozen tissue; reads were aligned to GRCh38 using the STAR two-pass workflow and gene-level quantification was performed as part of the GDC harmonization pipeline^9,62^. For IM151, RNA was extracted from macro-dissected FFPE tumor tissue using the High Pure FFPET RNA Isolation Kit (Roche), and whole-transcriptome libraries were generated using TruSeq RNA Access technology (Illumina)^27^. After removal of ribosomal reads, remaining reads were aligned to GRCh38 using GSNAP (version 2013-10-10), and gene-level read counts were quantified using the GenomicAlignments R/Bioconductor package and normalized to TPM^27^. For JR101, whole-transcriptome profiling was performed on FFPE tumor tissue (ACE version 3; Illumina NovaSeq) for 726 patients. Reads were aligned to the NCBI hs37d5 reference genome using STAR (version 2.4.2a-p1) and transcript-level values were generated by the Personalis ACE Cancer Transcriptome Analysis pipeline^5^. matrices provided by the original studies were log transformed log_2_(TPM+1) for downstream analysis. Genes were restricted to those measured in all three datasets. Within each dataset, genes were then ranked by median absolute deviation (MAD) across samples. The top 6,000 most variable genes according to the MAD were retained. The union of these per-dataset variable-gene sets yielded 7,762 genes used for downstream factorization.

#### Multi-dimensional NMF

Each dataset was factorized independently using the CoGAPS NMF algorithm with default parameters^30^, varying the factorization rank (K) from 10 to 25. For each run, the resulting gene-by-factor (amplitude) matrix was extracted and scaled such that each gene’s weights across factors summed to 1. CoGAPS gives us an Amplitude matrix A and Pattern matrix P such that:

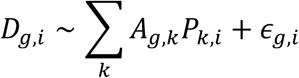

Where *_Ag, k_*is the weight of the *g*-th gene in the *k*-th factor, *_Pk,i_* is the pattern of the k-th factor in the i-th sample, and *_εg, i_* is residual error. If the *_Ag, k_* is scaled by ∑*_i Pk_*_,*i*_ each scaled factor reflects the total absolute gene expression of *g* associated with factor *k*

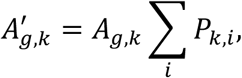

Normalizing each row of *_A_*^’^*_g,k_* gives us the fractional usage of g in the k-th factor

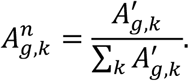

Scaled amplitude matrices from all runs and datasets were concatenated into a single combined matrix.

### Identifying consensus communities of factors

To identify recurrent factors, we computed a Pearson correlation matrix across factor gene-weight vectors and retained factors with correlation ≥0.6 to at least one other factor. Correlated factors were represented as a graph in which nodes corresponded to factors and edges were weighted by Pearson correlation. Graph layouts were generated using the Fruchterman–Reingold force-directed algorithm (ggraph v2.2.2 R package), and communities were identified using the Infomap algorithm on the weighted graph. Communities were retained if they contained factors from more than one dataset, with the minor dataset contributing at least 5% of factors within that community.

### Consensus Community Aggregation into Consensus Transcriptomic Programs

Within each community, genes were ranked for each factor by their scaled amplitude weights (highest to lowest across the 7,762-gene space), and subsequently aggregated into community associated gene weights using a modified Borda average where

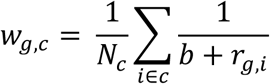

where *_Nc_* is the number of factors in the community and *_rg_*_,*i*_ is the rank of the *g*-th gene in the i-th factor based on the fractional allocation of the gene to the factor. b is a programmable buffer to avoid dominance of the top ranking over frequency of gene association. In this study, we use b=9. The top 100 genes were designated as marker genes and in conjunction with their associated aggregate weights defined the CTP. This approach is illustrated in **Figure S1**.

### Consensus Transcriptomic Program Scoring and Utilization

#### Calculating raw CTPscores

Samples were evaluated for each CTP using a weighted expression sum over the CTP marker genes. For each sample, we first computed a raw CTPscore **_CTP_***_c,i_* as the sum across marker genes of the product of (i) the aggregate CTP gene weight and (ii) the gene’s expression value log_2_(TPM+1) as

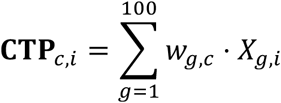

where *_wg,c_* is the weight of the g-th gene in the c-th community and *_Xg,i_* is the expression of the g-th gene in the i-th sample.

#### Calculating Normalized CTPscores

To account for the fact that weighted-sum statistics can be influenced by mean gene expression, we scaled each program’s raw statistic to an empirical null distribution generated by scoring gene-wise permuted data. Specifically, expression values were permuted independently for each gene by sampling across the input expression matrix (i.e., log_2_(TPM+1) of combined TCGA, IM151, and JR101 datasets), preserving each gene’s overall distribution while disrupting gene/gene relationships. This is illustrated in **Figure S2**. We generated a null dataset with 10× upsampling relative to the observed number of samples and recomputed the raw weighted-sum statistic to obtain a CTP-specific null distribution. The observed raw statistic was then standardized using the null mean and standard deviation to yield a z-score for each program; these z-scores are referred to throughout the manuscript as CTPscores.

#### CTP utilization analysis

CTP utilization was defined by converting each CTPscore to a one-sided p value (positive enrichment) and applying multiple-hypothesis correction across the 17 CTPs within each sample. Programs with sample-wise FDR ≤ 0.05 were considered “utilized,” indicating enrichment beyond that expected under the permutation-derived null. To assess patterns of co-utilization and mutual exclusivity among RCC-intrinsic programs, we constructed 2×2 contingency tables of binary utilization status for each pair of programs and performed Fisher’s exact test. Odds ratios greater than 1 indicated co-occurrence (programs utilized together more often than expected), while odds ratios less than 1 indicated mutual exclusivity. p values were corrected for multiple comparisons using the Benjamini-Hochberg method.

### Over-representation analysis

Over-representation analysis was performed on the 100 marker genes for each CTP using the fgsea R package with gene set collections from the Molecular Signatures Database (MSigDB), including Hallmark, Gene Ontology Biological Processes, and Positional (cytogenetic band) gene sets^63^. The top two enriched gene sets for each CTP within each collection are visualized in **Figure 2A**.

### Inferring cellular specificity of consensus transcriptomic programs

Processed scRNAseq count matrices and accompanying cell annotations were obtained from the published datasets (Li et al.,^35^ Zhang and Narayanan et al.,^36^ and Alchahin and Mei et al.^37^) and analyzed in R using Seurat (v5.3.1). To enable consistent cross-study comparisons, original author-provided cell labels were harmonized into seven consolidated compartments (RCC, B-cell, T/NK cell, Stromal, Endothelial, Myeloid/APC, and benign epithelium; **Table S3**). Within each scRNAseq dataset, raw counts were normalized and variance-stabilized using SCTransform, using the SCTransform corrected counts as the expression input for subsequent analyses. Principal component analysis (PCA) was performed on SCTransform-transformed features and visualized using Uniform Manifold Approximation and Projection (UMAP) based on the same PC space.

#### From scRNAseq

CTP scoring and utilization in scRNAseq data were computed using the rC3TP package as described above, using SCTransform corrected counts as input. To assess cell-type specificity of each program, we constructed, for each dataset and each CTP, a 2×2 contingency table comparing (i) cells of a given harmonized cell-type category versus all other cells and (ii) program utilization versus non-utilization. Odds ratios and p values were computed using Fisher’s exact test within each dataset, with multiple testing correction applied across tested programs and cell-type categories.

#### From RCC Tumorgrafts

RCC tumorgraft log-transformed TPM expression matrices and sample annotations were obtained from the supplementary materials of Elias et al.^38^ CTPscores and utilization were computed using the rC3TP package. Because the tumorgraft cohort is modest in size and includes programs whose marker genes can have low absolute expression in this setting, we referenced utilization testing to the empirical null distribution derived from the merged bulk RCC cohorts (TCGA, IMmotion151, and JAVELIN Renal 101), rather than estimating a tumorgraft-specific null. This approach reduces spurious “utilization” calls that can arise when within-dataset null variance is artificially small for lowly expressed marker sets (i.e., non-malignant genes).

### Association of CTPs with genomic alterations and fusions

Somatic mutation data were obtained for each of the three discovery cohorts, which differed in sequencing platform and variant calling approach. For the TCGA, matched tumor-normal whole exome sequencing (WES) was performed on fresh-frozen tissue, and somatic variant calls were downloaded from the GDC^62^, which aligns reads to GRCh38 and applies four independent variant calling pipelines (MuTect2, MuSE, VarScan2, and Pindel) with germline filtering. For JR101, tumor-only WES was performed on FFPE tissue using the Personalis ACE Cancer Exome pipeline, which employs proxy-normal filtering to remove germline variants. For IM151, tumor-only comprehensive genomic profiling was performed on FFPE tissue using the FoundationOne assay. Mutation calls for JR101 and IM151 were accessed from deposited data accompanying the original publications. These deposited data also included *TFE3*/*TFEB* fusion calls and *CDKN2A* deletion calls for the IM151 dataset. *TFE3/TFEB* fusion status for the TCGA dataset was accessed from deposited supplementary material in Bakouny et al.^60^ *CDKN2A* deletion status was obtained from the GDC gene level copy number alteration pipeline^62^. Full details of DNA extraction, library preparation, sequencing, and variant calling for each cohort are described in the respective primary publications^5,27,62^.

To test associations between CTPscores and canonical driver alterations, we performed logistic regression within each dataset independently. Alterations tested included somatic mutations in *VHL, BAP1, PBRM1, SETD2, TSC1, TSC2, PTEN, MTOR, PIK3CA, KDM5C, NF2, FAT1*, and *TP53*, as well as *CDKN2A* deletions and *TFE3/TFEB* fusions where available. Alteration status was coded as a binary outcome (1 = altered, 0 = wild-type), and each CTPscore was used as the predictor variable.

Regression coefficients, confidence intervals, and p values were extracted for each CTP-gene pair. p values were corrected for multiple comparisons across all tests within each dataset using the Benjamini-Hochberg method. Select associations were visualized using box plots comparing altered versus wild-type cases, and receiver operating characteristic (ROC) curves were generated for a subset of CTP–mutation pairs.

### Trajectory inference and transcriptomic stage assignment

Prior methods have applied pseudotime to bulk gene expression directly^64^, here we instead use RCC-intrinsic CTPscores as input, ensuring the trajectory space is restricted to tumor cell states without requiring a priori covariate specification. To model transcriptomic progression among RCC-intrinsic programs, we performed diffusion map analysis using the destiny R package. The input matrix consisted of CTPscores for five RCC-intrinsic programs (R1, R2, R3, R4, and MP-Prolif), selected based on their association with canonical ccRCC driver mutations. The analysis was restricted to 1,463 tumor specimens (70.0%) that significantly utilized at least one VHL-associated program (R1–R4). A diffusion map was computed with k = 300 nearest neighbors and 20 eigenvectors, and diffusion pseudotime (DPT) was calculated from the resulting map.

Diffusion components (DCs) were visualized in pairwise scatter plots. The first three components (DC1, DC2, DC3) were retained for trajectory analysis, with DPT used as the operational pseudotime metric for downstream analyses.

Assignment of trajectory and transcriptomic stage (TS) is implemented in the rC3TP R package (this publication), which provides two modes: a stringent mode, which assigns trajectory and TS only to samples that significantly utilize at least one of R1-R4 or MP-Prolif, and an inclusive mode, which assigns trajectory and TS to all samples based on the maximum CTPscore or CTPscore ratio regardless of utilization significance.

Samples that do not meet the utilization threshold in stringent mode are labeled as “Other.” Stringent mode was used for all analyses unless otherwise specified.

Trajectory was assigned based on the relative magnitude of R2 and R4 CTPscores: samples with R4 > R2 were classified as *BAP1*-like, and those with R2 > R4 as *PBRM1*-like. TS was assigned based on the dominant RCC-intrinsic CTP for each sample: R1-dominant tumors were classified as Early, R2- or R4-dominant as Intermediate, and R3-or MP-Prolif-dominant as Late.

To assess the relationship between TS and the tumor microenvironment, TME CTPscores (TME-Endo, TME-Tcell, TME-Myelo, TME-Stroma) were compared across CTP groups stratified by trajectory using Wilcoxon rank-sum tests, and Spearman correlation was used to test for monotonic trends across the ordered TS stages. The association between DPT and nuclear grade was assessed in TCGA using pairwise Wilcoxon tests across grade categories (G1, G2, G3, G4).

### Spatial transcriptomics

Processed Visium count matrices from eight sarcomatoid ccRCC cases were accessed from the supplementary material of Salgia et al.^51^. Histological annotations assigning each spot to a tissue compartment (clear cell [CC], sarcomatoid [Sarc], mixed, stroma, or necrosis) were provided per the original publication. Count data were processed using Seurat, and CTPscores were calculated at the spot level on SCTransform-normalized counts using the approach detailed above, as implemented in the rC3TP package. Trajectory and TS were assigned using the inclusive mode, as spot-level UMI counts varied substantially across specimens and influenced utilization calls under the stringent threshold. Spot-level CTPscores, TS, and trajectory assignments were projected onto tissue coordinates using Seurat’s spatial visualization functions and overlaid with matched H&E images. To quantify spatial patterns of trajectory co-occurrence, we compared CTPscores at the spot level using scatter plots of utilization status, and calculated odds ratios using Fisher’s exact tests. The distribution of TS across histological compartments (CC vs. Sarc) was compared using chi-squared tests.

### Multiregional sequencing

Bulk RNAseq TPM matrices and matched somatic mutation calls from the TRACERx Renal cohort (213 regions from 64 patients) were accessed from supplementary material of Fernández-Sanromán et al.^41^ TPM values were log₂-transformed, and region-level CTPscores, trajectory, and TS were assigned using the rC3TP package with default (stringent) settings. The number of somatic alterations and whole-genome instability index (WGII) per region were obtained from the published supplementary material. To visualize intra-tumoral progression, CTPscores for R1-R4 and MP-Prolif were displayed alongside driver mutation status in heatmaps organized by unsupervised hierarchical clustering. Patients were classified as TS-concordant if all sampled regions were assigned to the same stage (Early/Intermediate or Late), and TS-discordant if at least one region was classified as Late and another as Early or Intermediate; regions classified as “Other” were excluded from concordance assessment. Intra-patient trajectory concordance was quantified as the fraction of region pairs sharing the same trajectory assignment and compared to a null distribution generated by permutation. The number of somatic alterations and WGII were compared between Early/Intermediate and Late regions using Wilcoxon rank-sum tests.

### Clinical Outcomes Analysis

Clinical data for the KIRC subset of the TCGA were accessed through the Clinical Data Resource^65^ and included overall survival (OS), defined as time from specimen collection to death, with a median duration of follow-up of 55.1 months (IQR 32.1–81.7). The JR101 and IM151 cohorts are phase III clinical trials in the front-line metastatic setting comparing avelumab plus axitinib (JR101) and atezolizumab plus bevacizumab (IM151) to sunitinib^5,27^. All patients in these cohorts had metastatic disease at the time of study enrollment. These trials reported progression-free survival (PFS), defined as time from initiation of treatment to disease progression or death. Median duration of follow-up was 16.6 months (IQR 13.8–20.5) in the IM151 cohort and 11.0 months (IQR 7.0–15.2) in the JR101 cohort.

Univariate associations between clinical outcomes and individual CTP utilization, trajectory assignment, and TS were assessed using Cox proportional hazards regression within each dataset. Hazard ratios (HRs) and 95% confidence intervals were estimated for each predictor and visualized using forest plots. Kaplan-Meier survival curves were generated for TS categories (Early, Intermediate, Late, and Other) and compared using the log-rank test.

Multivariable Cox regression was performed to assess the independent prognostic value of TS. In the TCGA, models were adjusted for pathologic stage (I–IV) and nuclear grade (G1/G2, G3, G4). In IM151, models were adjusted for treatment arm (atezolizumab plus bevacizumab vs. sunitinib) and IMDC risk score (favorable, intermediate, poor). In JR101, models were adjusted for treatment arm (avelumab plus axitinib vs. sunitinib). A separate multivariable model was performed for the subset of specimens with available genomic data, additionally including a *BAP1/PBRM1* mutation classifier as a covariate, categorized as *BAP1* mutant, *PBRM1* mutant, double mutant (*BAP1* and *PBRM1*), or wild-type for both, as previously described^42,43^.

To test whether TS or individual CTPs predicted differential treatment benefit, interaction analyses were performed in IM151 and JR101. Patients were stratified by CTP utilization status (utilized vs. not utilized) and treatment arm, and PFS was compared between treatment arms within each stratum using log-rank tests.

## QUANTIFICATION AND STATISTICAL ANALYSIS

Specific statistical details, sample size, and baseline characteristics, for every figure and analysis are reported in the corresponding figure legend, STAR methods section, supplementary table, and explicitly commented on in the deposited analysis code. All analyses were performed in R (v4.5.1) and Rstudio unless otherwise specified.

Inclusion and exclusion criteria for samples and clinical groupings are described in the relevant STAR Methods sections. For all figures, *p < 0.05, **p < 0.01, ***p < 0.001, unless otherwise noted in the figure legend. Unless otherwise noted, p values were adjusted for multiple comparisons using the Benjamini-Hochberg method throughout the manuscript.

For all box plots, the centerline represents the median, box limits represent the upper and lower quartiles (25th and 75th percentiles), and whiskers extend to the most extreme values within 1.5 times the interquartile range. Individual data points are overlaid where indicated. Violin plots display the full distribution of data with the same median and quartile conventions. Kaplan-Meier survival curves were compared using the log-rank test, and numbers at risk are displayed below each plot.

Heatmaps were generated using the ComplexHeatmap R package, with unsupervised hierarchical clustering performed using Euclidean distance and Ward.D linkage unless otherwise specified. Dot plots, box plots, violin plots, bar plots, scatter plots, forest plots, and histograms were generated using ggplot2. UMAP embeddings for scRNAseq data and spatial maps for Visium data were generated using Seurat (v5.3.1) plotting functions.

## SUPPLEMENTARY FIGURES

**Figure S1.**
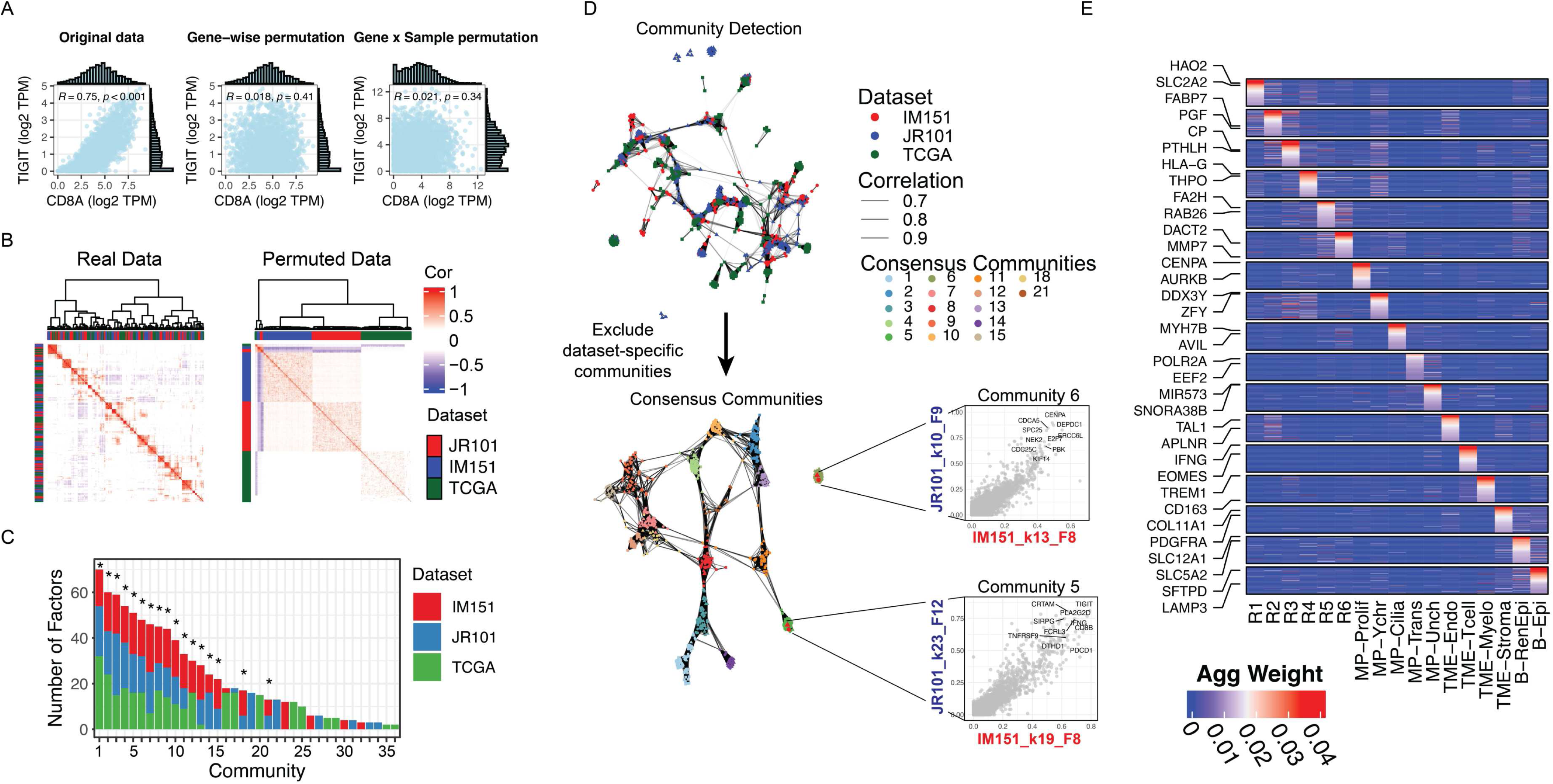
A subset of NMF factors are recurrent across RCC datasets (Related to Figure 1). (A) Representative scatter plot of log_2_(TPM+1) for two correlated genes, *CD8A* and *TIGIT* demonstrating observed data (left), gene-wise permuted data (center), and gene by sample permutation (right). Histograms across the x- and y-axis reflect the distribution of the *CD8A* and *TIGIT* gene, respectively. (B) Heatmap of pairwise Pearson correlations among all scaled NMF factors generated across datasets (840 total factors). Factors are organized by unsupervised clustering, with top and left annotations indicating dataset of origin. The left heatmap (observed data) reveals cross-dataset clusters of correlated factors that are absent in the row-wise permuted control (right). (C) Graph-based representation of correlated factors where edges correspond to Pearson r and nodes correspond to individual NMF factors (Top). Communities containing factors from 2 or more datasets were considered “consensus” communities and retained (bottom). Inset scatter plots demonstrate representative cross-dataset correlations of NMF factors belonging to a consensus community. Values reflect scaled NMF factor gene weights. (D) Stacked bar plot summarizing the number of factors per community and their dataset of origin. Communities comprised of a mixture of datasets (at least 5% from minor dataset) are considered “consensus” and marked with an asterisk (*). (E) Heatmap of aggregate CTP marker gene weights (methods).

**Figure S2.**
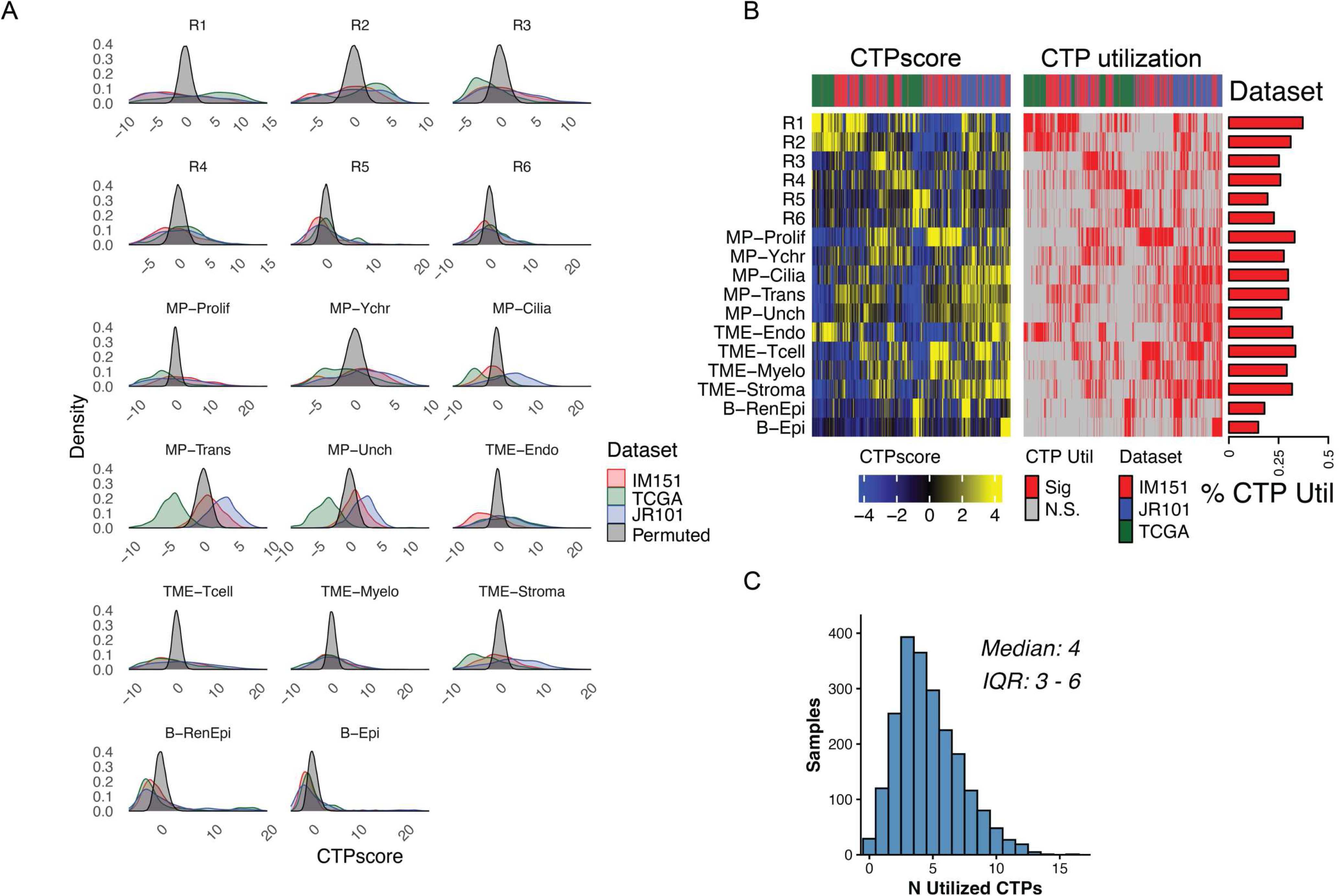
Defining CTPscore and utilization (Related to Figure 1). (A) Density plot depicting distribution of CTPscores in gene-wise permuted data (null; black), IM151 (red), TCGA (green), and JR101 (blue) datasets. The x-axis depicts a z-transformation based on mean and standard deviation of the null distribution. (B) Heatmap of CTPscores (left) and CTP utilization (right) across the TCGA, IM151, and JR101 datasets. Bar plot annotation depicts the fraction of samples utilizing a given CTP. (C) Distribution of CTP utilization across the combined cohort (n = 2,163). Utilization is defined as a CTPscore significantly exceeding a gene-wise permuted null distribution (FDR < 0.05). The histogram displays the number of concurrent utilized programs per sample (median = 4; Interquartile range [IQR] 3-6).

**Figure S3.**
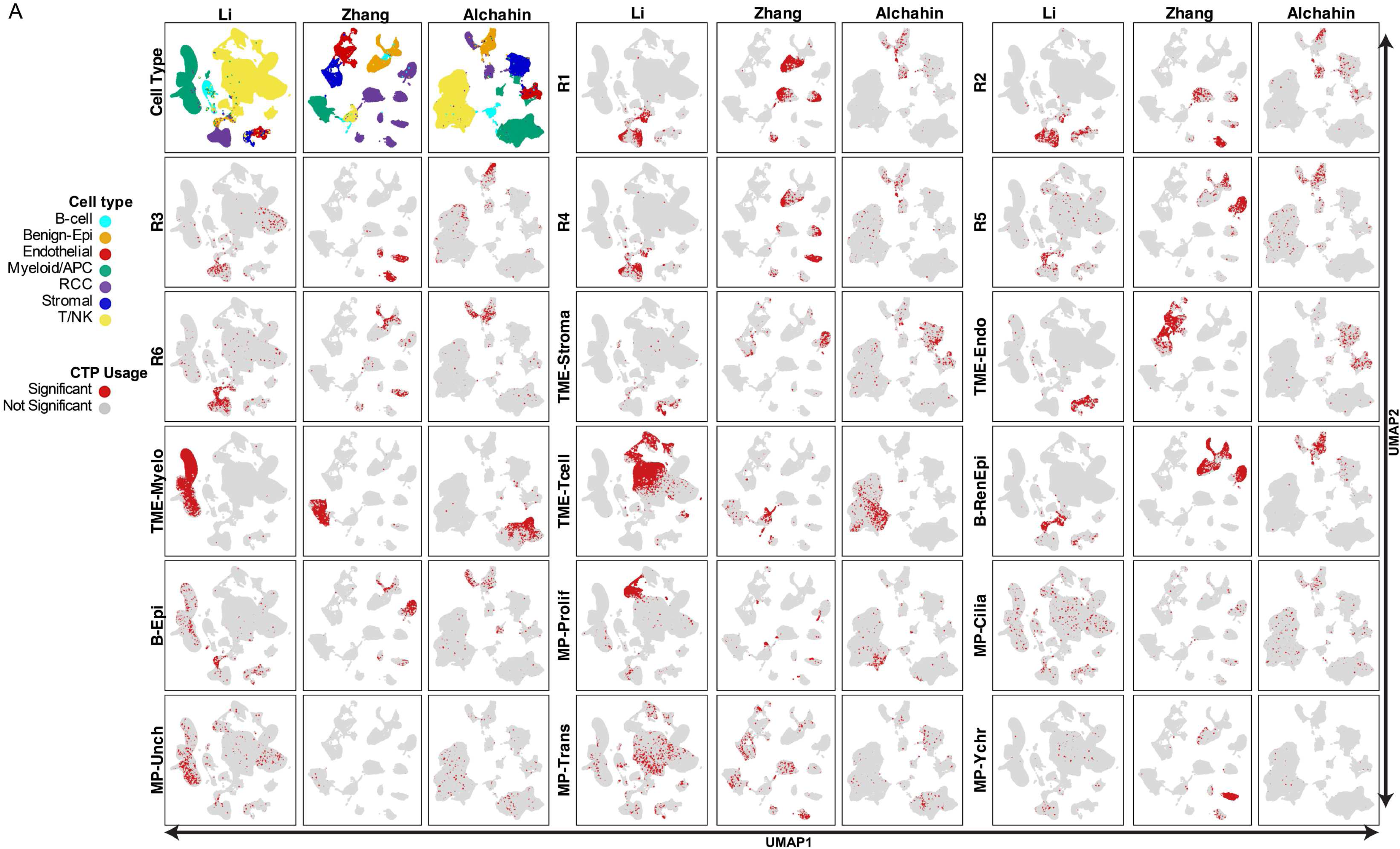
CTP distribution across RCC scRNAseq datasets (Related to Figure 2). UMAP embeddings of three RCC scRNAseq cohorts^35–37^ highlighting cells with statistically significant utilization of each CTP (n = 17 CTPs).

**Figure S4.**
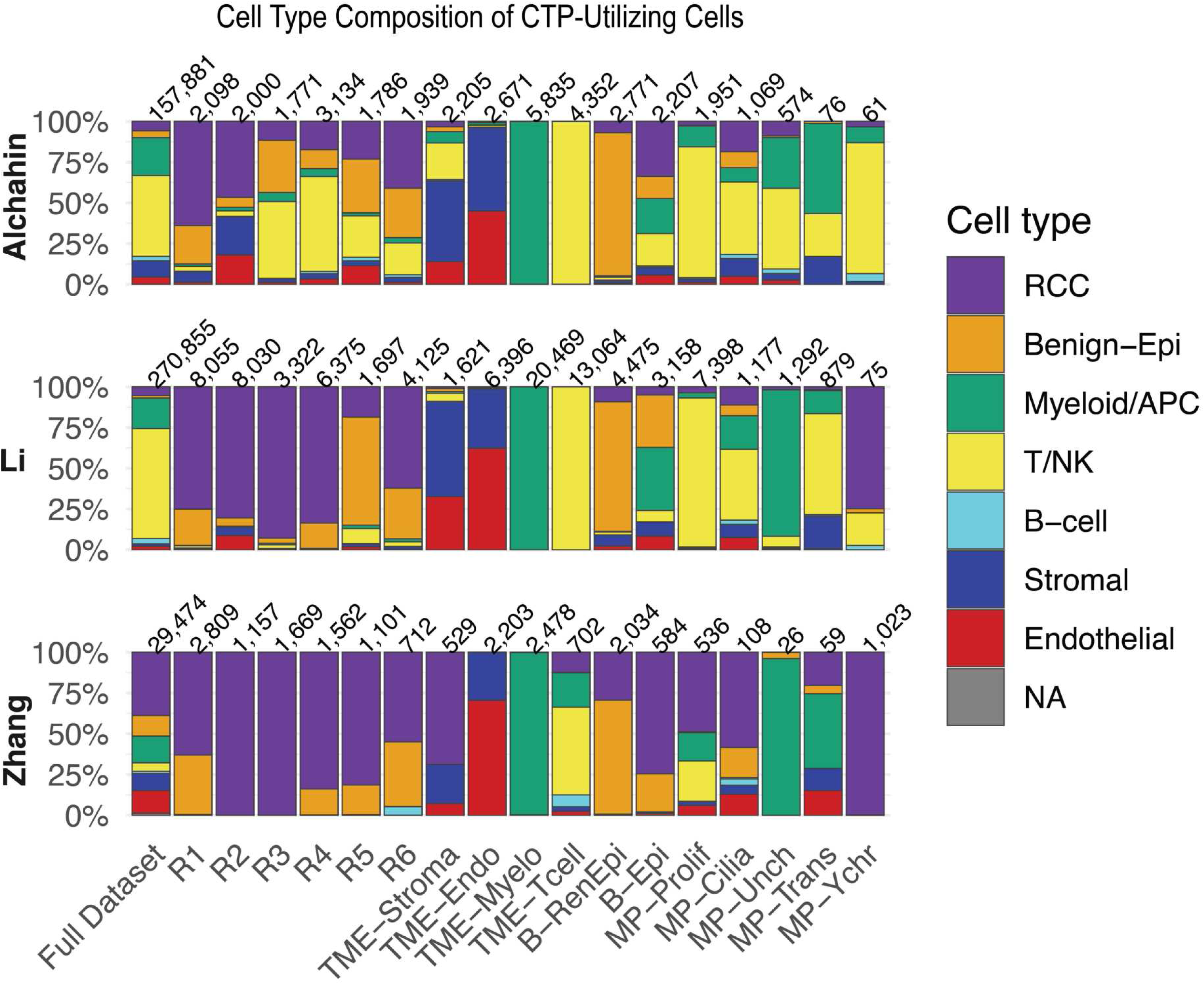
Composition of CTP-utilizing cells across RCC scRNAseq datasets (Related to Figure 2). Stacked bar plot showing the proportion of cells utilizing each CTP, stratified by dataset. The first column depicts the composition of the full dataset. Numbers at the top of the stacked bar plot depict the number of cells utilizing a given CTP.

**Figure S5.**
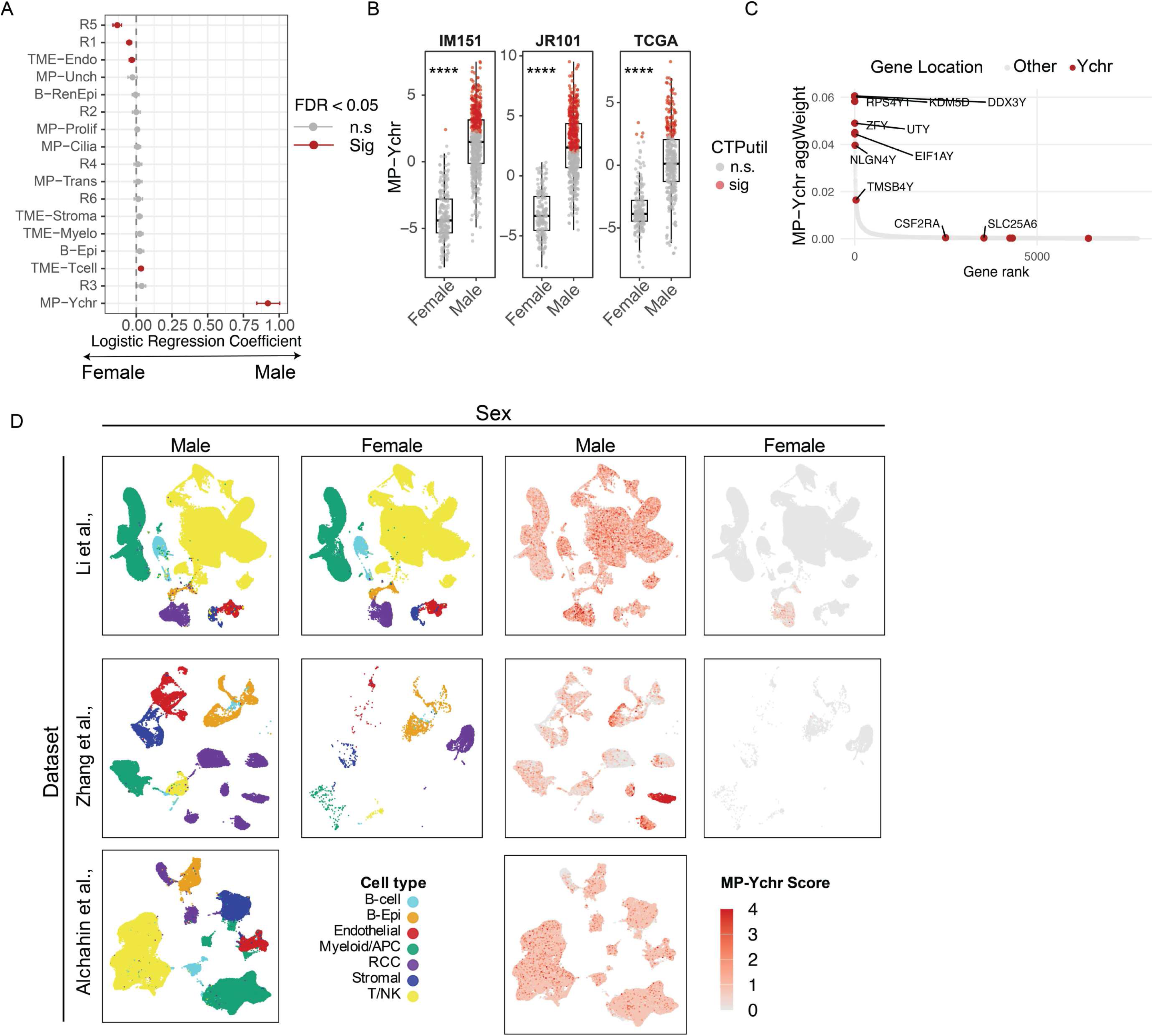
MP-Ychr is a sex-linked CTP (Related to Figure 2). (A) Forest plot of logistic regression coefficients (±95% CI) testing the association between CTP utilization and sex across CTPs in bulk RNAseq cohorts (TCGA, JR101, IM151). (B) Boxplots of MP-Ychr scores stratified by sex and dataset; color indicates MP-Ychr utilization. (C) MP-Ychr marker genes ranked by their aggregate weight; genes encoded on the Y chromosome are highlighted in red. (D) Distribution of MP-Ychr scores in scRNAseq datasets stratified by sex; note that the Alchahin cohort was comprised of solely male subjects.

**Figure S6.**
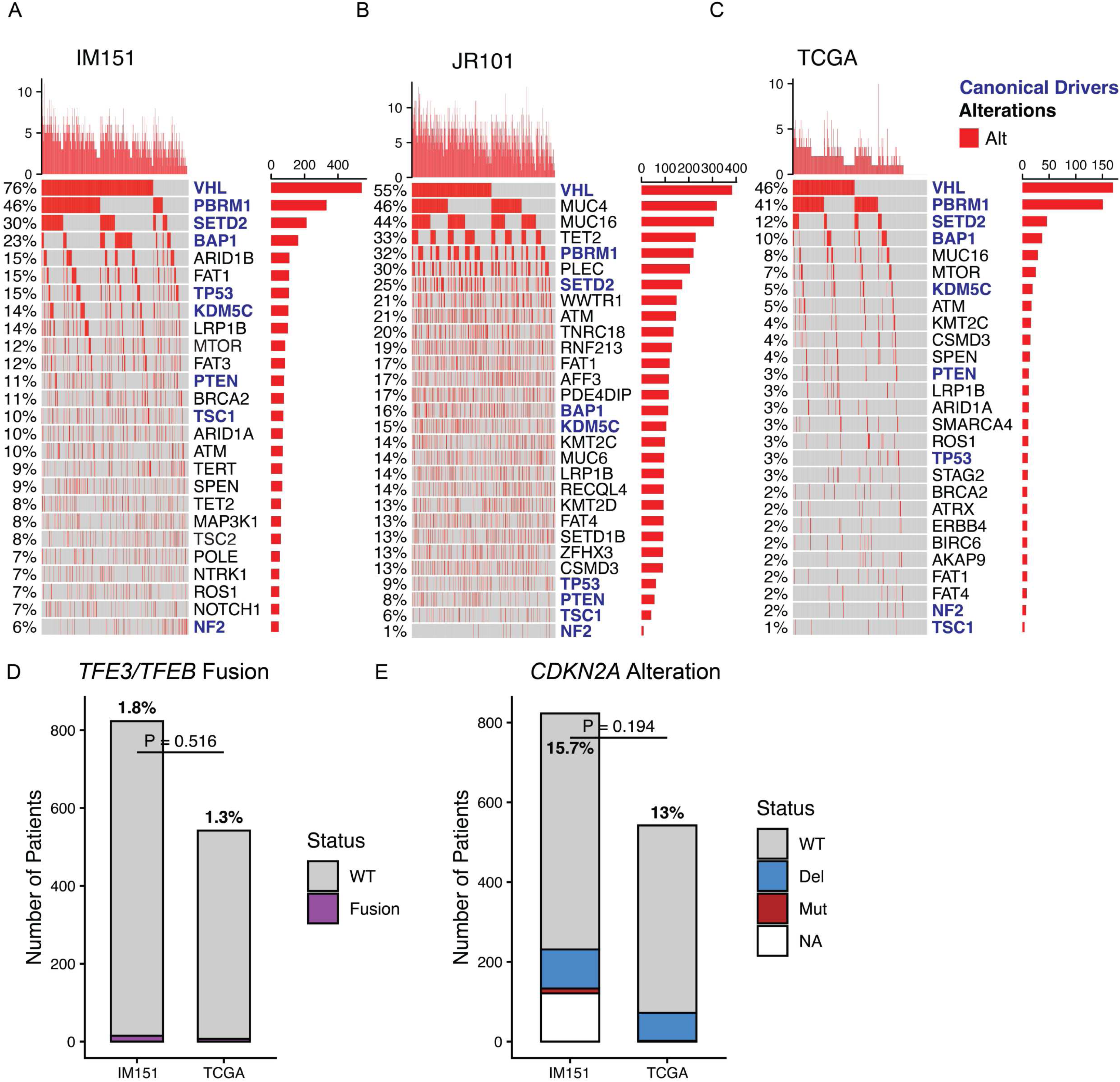
Mutation landscape across the TCGA, IM151, and JR101 datasets (Related to Figure 3). (A)-(C) Oncoprint of most frequent mutated genes and canonical ccRCC driver genes (blue) in the (A) TCGA, (B) IM151, and (C) JR101 datasets. (D) Frequency of *TFE3* or *TFEB* fusions and (E) *CDKN2A* deletions in the TCGA and IM151 datasets where available.

**Figure S7.**
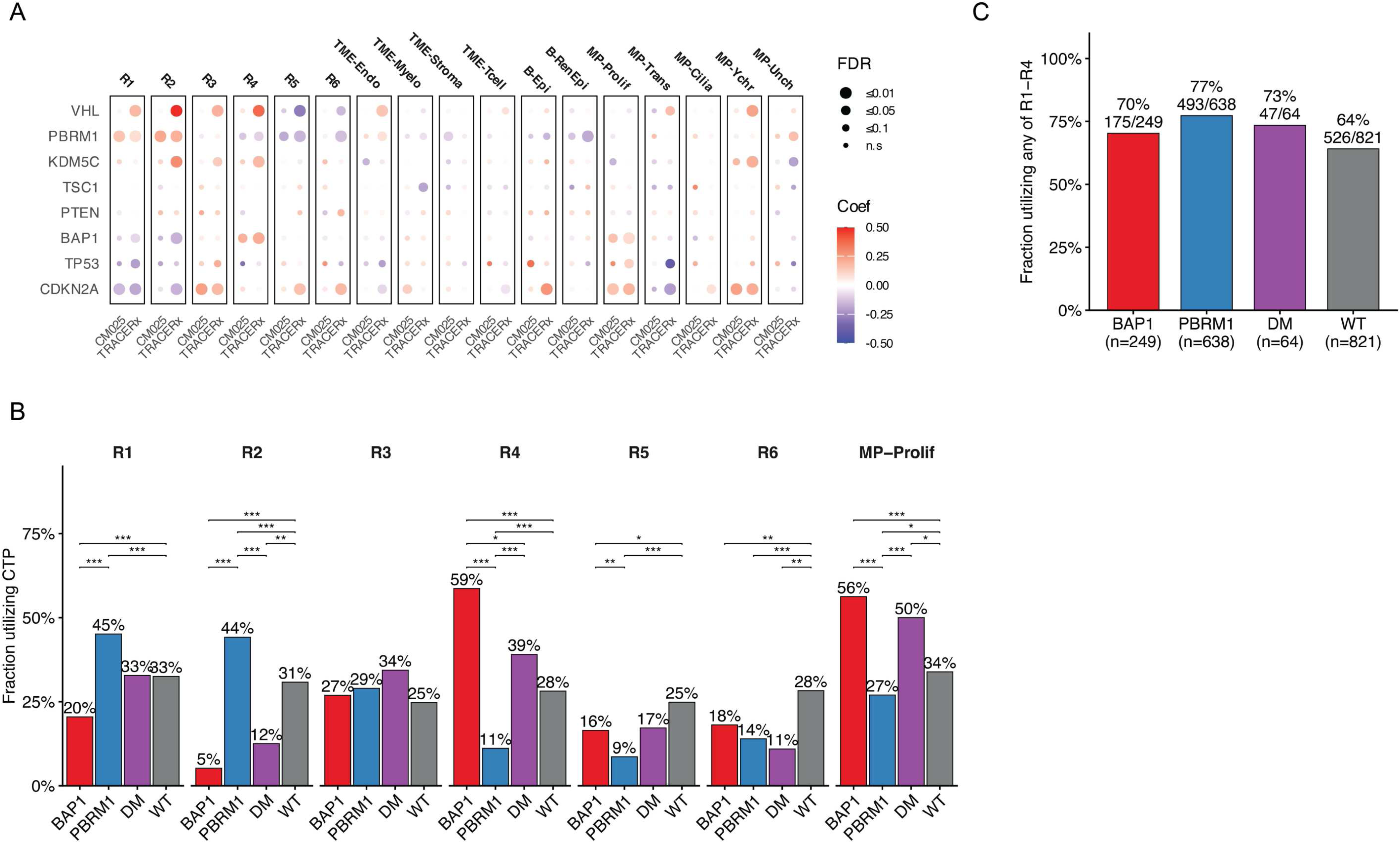
Validation of CTP–driver associations and BAP1/PBRM1 stratification of CTP utilization (Related to Figure 3). (A) Dot plot displaying the association between CTPscores and canonical driver alterations across the Checkmate 025/010/009^17^ (N = 217) and TRACERx^41^ (N=230) cohorts. Associations were assessed via logistic regression within each dataset independently. Dot size represents statistical significance (−log₁₀ FDR), and dot color represents the regression coefficient (red, positive association; blue, negative association). (B) Bar plot of CTP utilization stratified by BAP1/PBRM1 mutation status across the discovery cohorts (TCGA + IM151 + JR101; with mutation data, n=1,772). Each facet shows the percentage of patients in a given mutation group (BAP1, n=249; PBRM1, n=638; double mutant [DM], n=64; wild-type [WT], n=821) whose tumor utilizes the indicated CTP. Brackets indicate significant pairwise comparisons (Fisher’s exact test, BH-adjusted across all 42 contrasts; *p≤0.05, **p≤0.01, ***p≤0.001). (C) Fraction of cases in each *BAP1/PBRM1* group whose tumor utilizes at least one of R1–R4. Values above each bar give the percent utilization and the count of utilizing cases over the group total.

**Figure S8.**
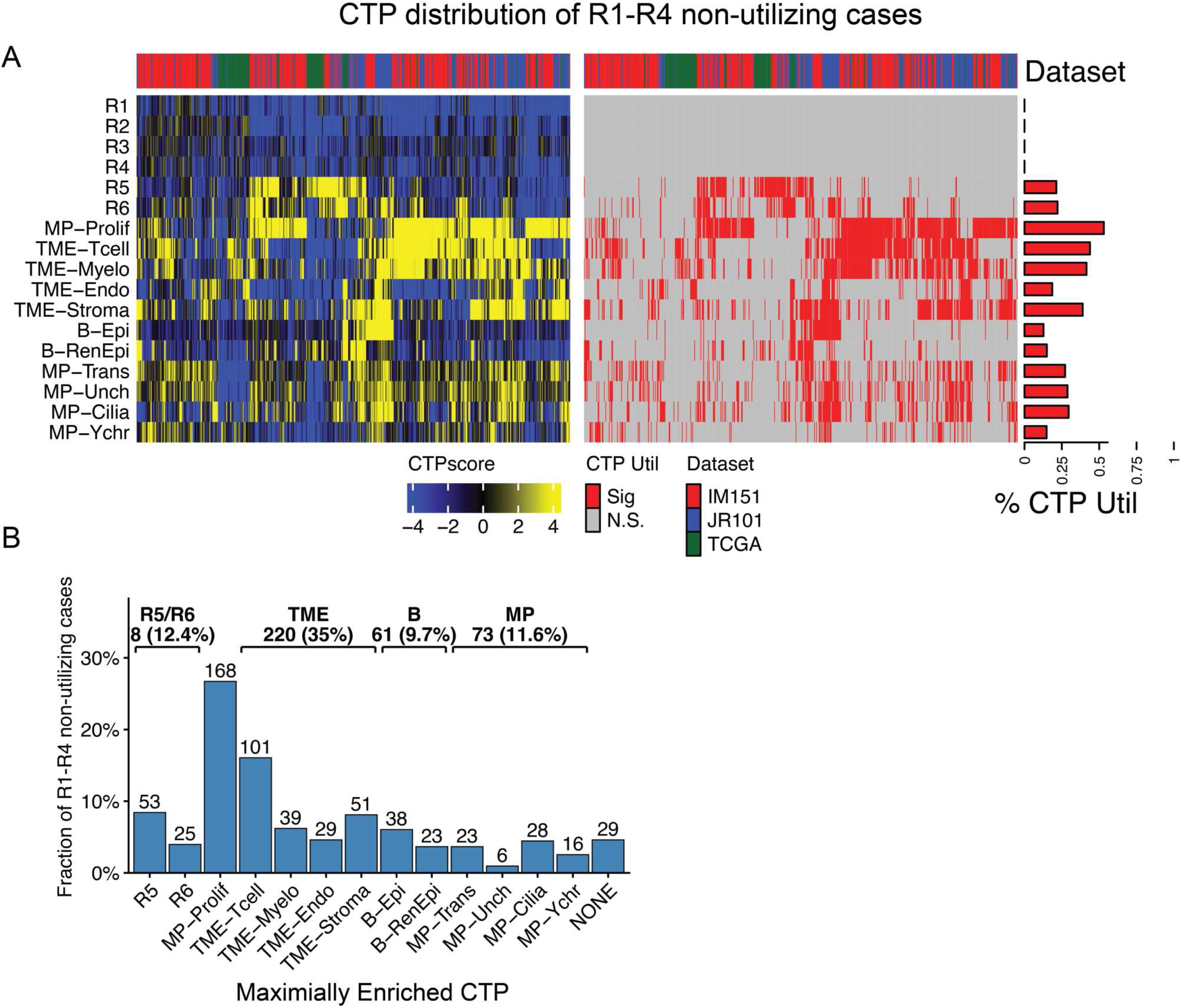
CTP distribution in R1–R4 non-utilizing cases (Related to Figure 4). (A) Combined CTPscore and utilization heatmaps across all 17 CTPs for unified-cohort tumors that do not utilize R1–R4 (n = 628 of 2,091; 30%). Utilization is defined as a CTPscore significantly exceeding a gene-wise permuted null (FDR < 0.05). Columns are jointly clustered; top annotation = dataset, right annotation = per-CTP utilization fraction. (B) CTPmax distribution across the same 628 tumors; CTPmax is the CTP with the highest score among utilized CTPs (“NONE” if none utilized). Brackets summarize R5/R6 (n = 78), TME (n = 220), benign (n = 61), and MP (n = 73) groupings.

**Figure S9.**
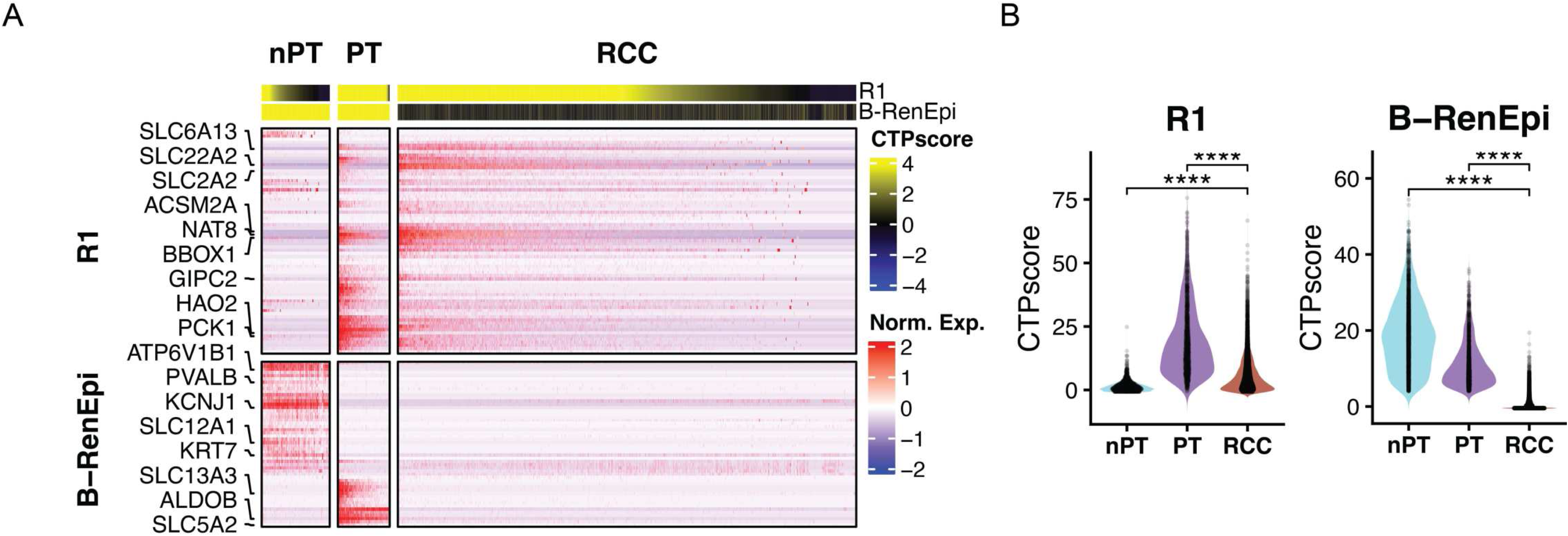
R1 captures a proximal tubule lineage program which persists in early ccRCC (Related to Figure 4). (A) Heatmap of R1 and B-RenEpi scores and selected marker gene expression across proximal tubule (PT), non–proximal tubule epithelium (nPT), and malignant RCC cells in the Li et al. scRNAseq cohort. Marker genes were filtered to those with non-zero expression in ≥1% of cells. (B) Violin plots of R1 and B-RenEpi scores stratified by cell type. p values were calculated using the Kruskal–Wallis test.

**Figure S10.**
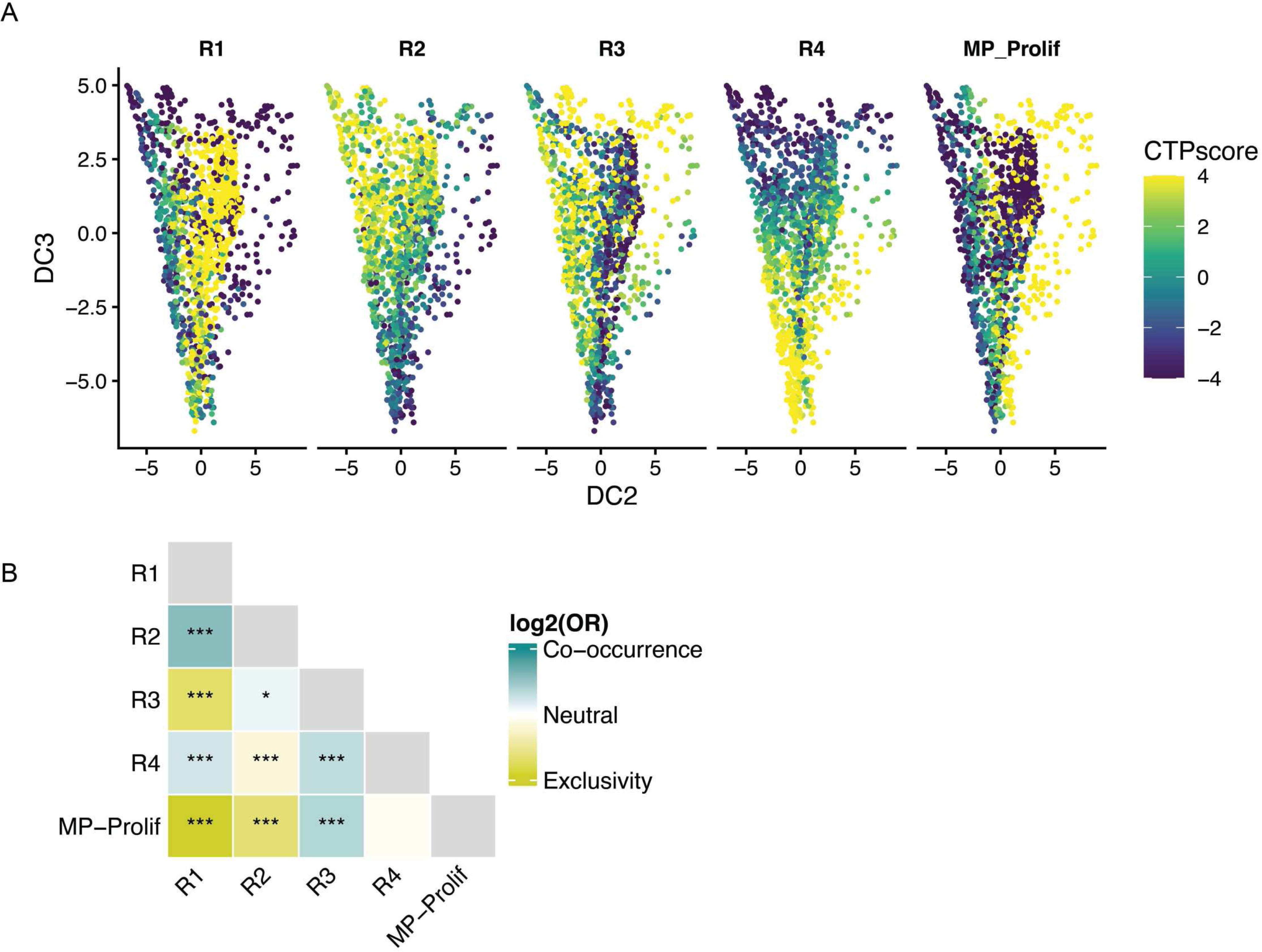
Diffusion mapping and co-utilization patterns support an R2–R4 branch point (Related to Figure 4). (A) Scatter plot of diffusion components DC2 and DC3, faceted by R1–R4 and MP-Prolif CTPscores. (B) Heatmap summarizing pairwise co-utilization versus mutual exclusivity among R1–R4 and MP-Prolif across the combined TCGA, JR101, and IM151 cohorts. Odds ratios and p values were calculated using Fisher’s exact test.

**Figure S11.**
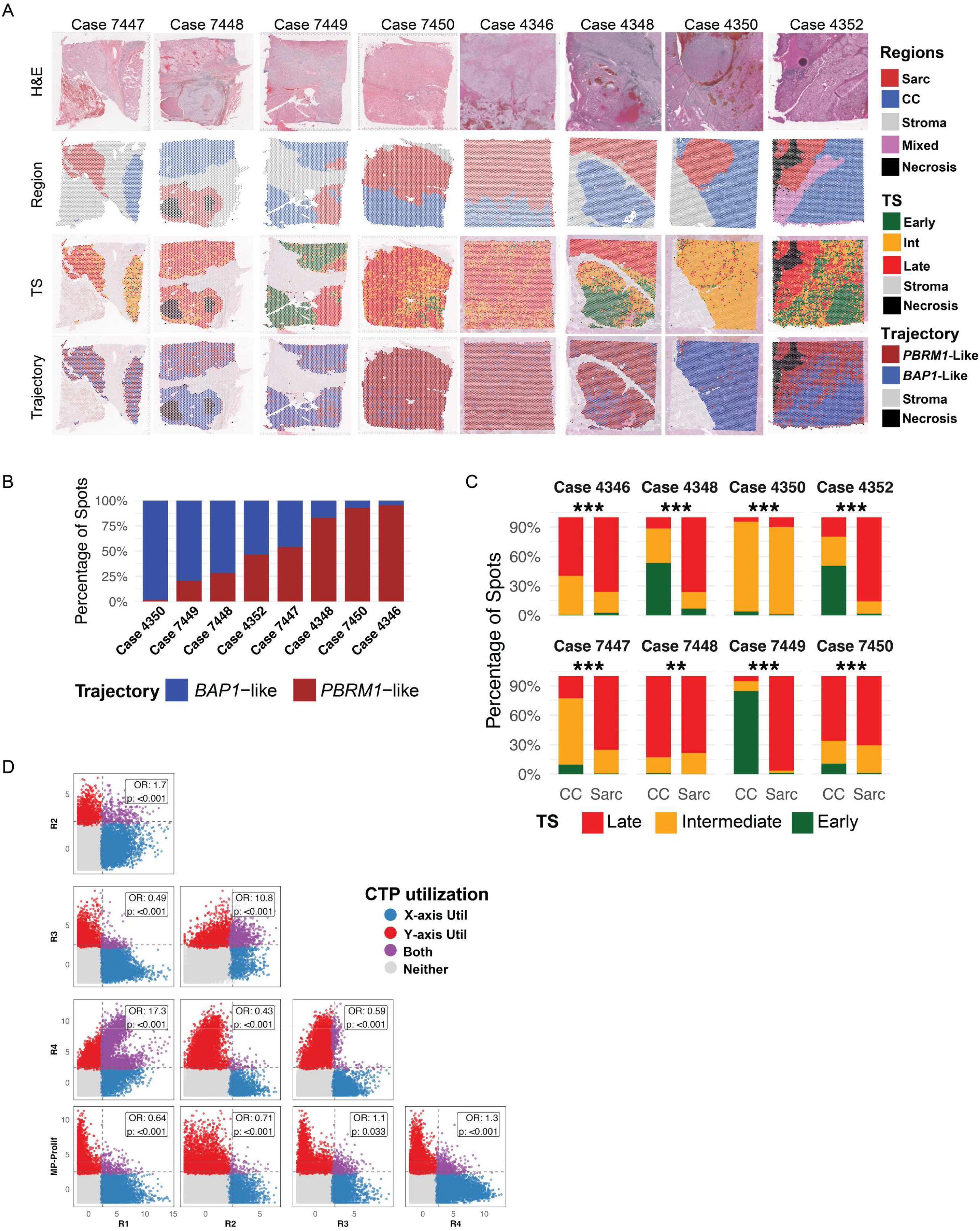
Spatial transcriptomics resolves trajectory and transcriptomic stage at spot-level resolution (Related to Figure 5). (A) Spatial maps showing H&E, published region annotations (Salgia et al.), spot-level transcriptomic stage, and spot-level trajectory assignment. (B) Stacked bar plot showing the fraction of spots per patient assigned to *BAP1*-like versus *PBRM1*-like trajectories. (C) Stacked bar plot comparing the distribution of transcriptomic stage across spots in sarcomatoid (Sarc) versus conventional clear cell (CC) regions. (D) Scatter plots showing spot-level CTPscores and utilization for R1–R4 and MP-Prolif. Odds ratios and p values were calculated using Fisher’s exact test.

**Figure S12.**
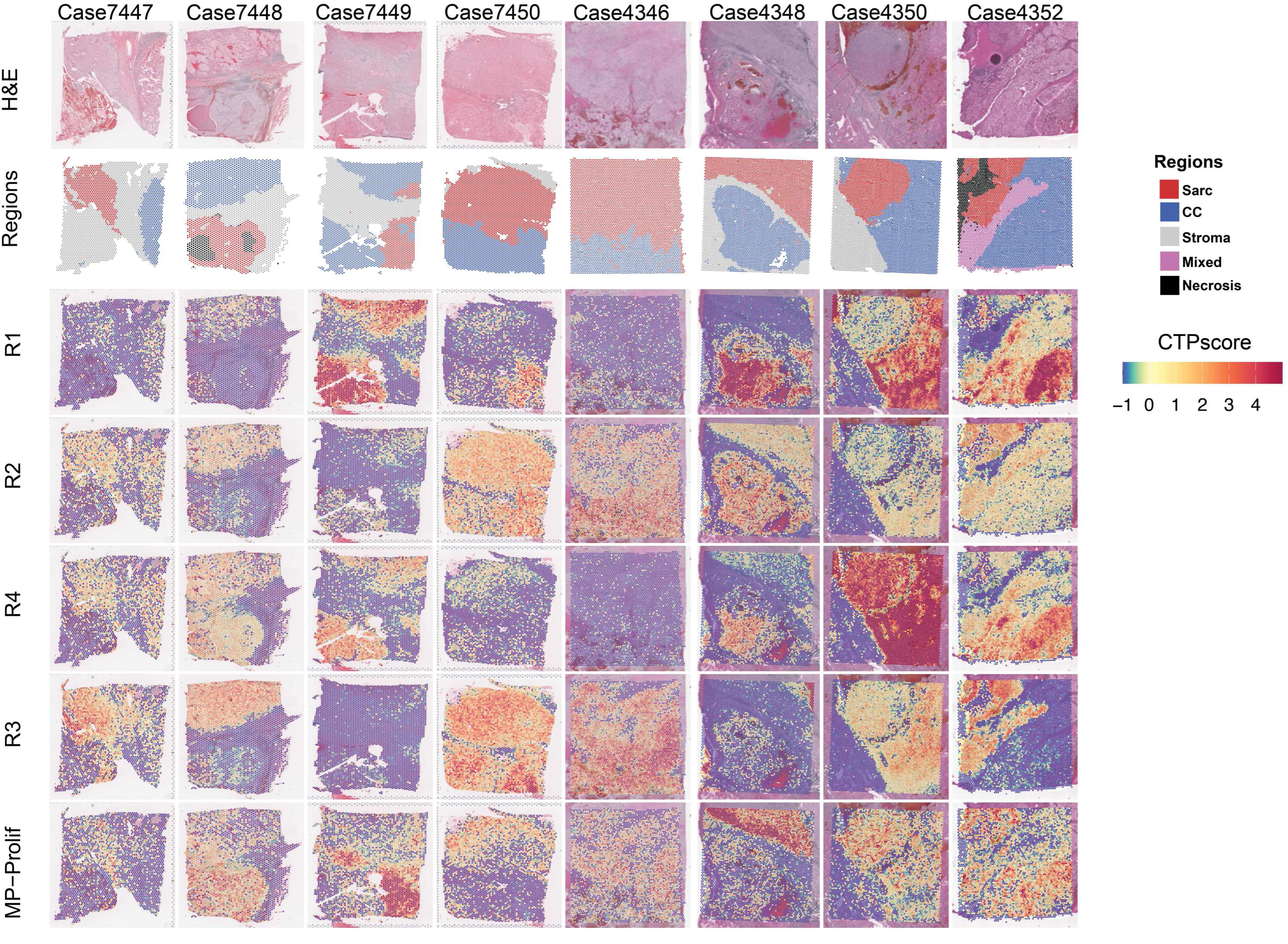
Spatial distribution of transcriptomic stage associated CTPs (Related to Figure 5). (A) Spatial maps from the Salgia et al. cohort showing H&E, published region annotations, and spot-level CTPscores for the RCC-intrinsic programs R1–R4 and MP-Prolif.

**Figure S13.**
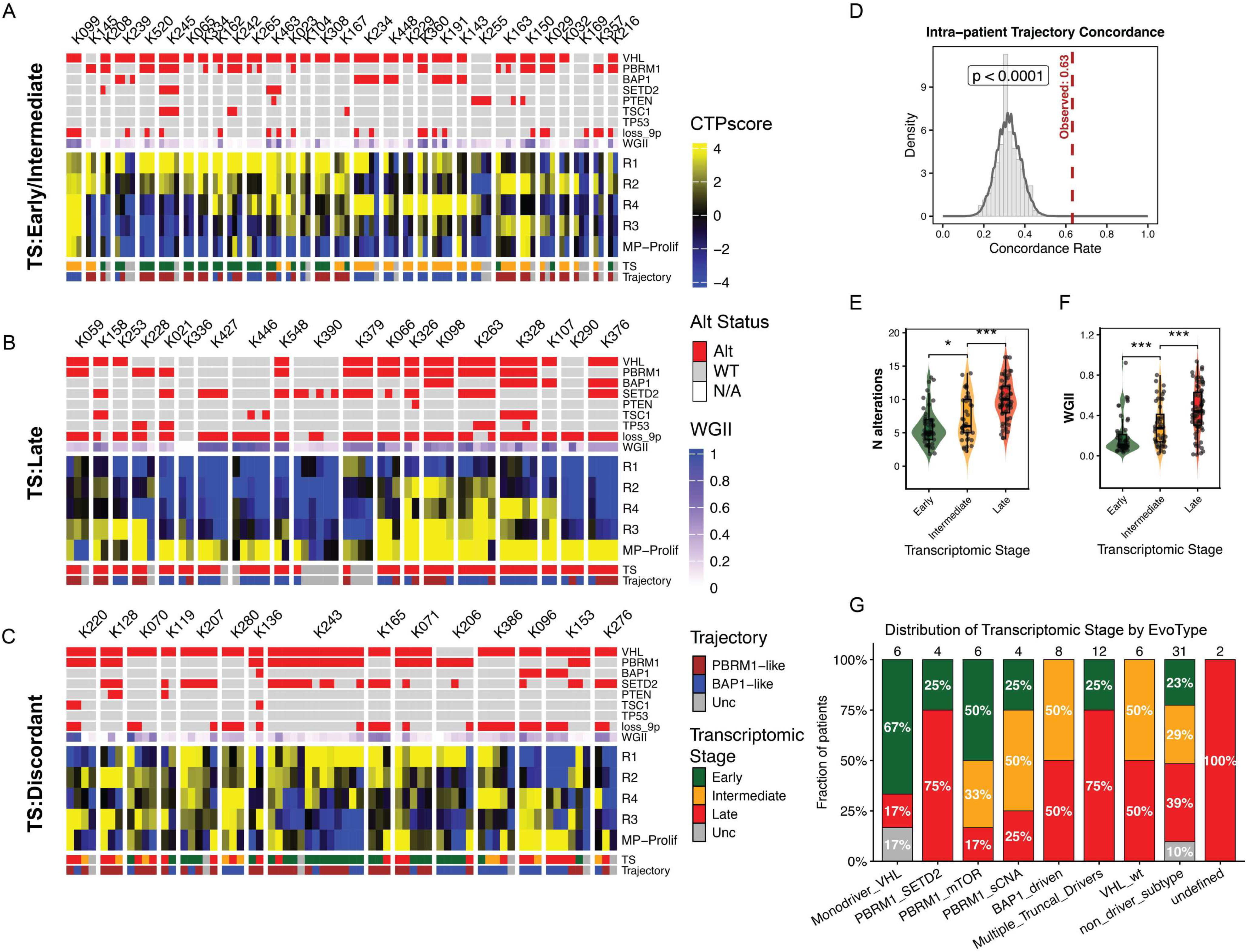
CTP distributions across multi-regional sequencing in the TRACERx cohort (Related to Figure 5). (A–C) Combined oncoprint and heatmap summarizing region-level driver alterations, transcriptomic stage (TS), trajectory assignment, and CTPscores (R1–R4 and MP-Prolif) across multi-regional samples. Patients are shown with: (A) Early/Intermediate TS only, (B) Late TS only, or (C) discordant TS across sampled regions. Columns represent individual regions and are grouped by patient to highlight within-patient heterogeneity. (D) Intra-patient trajectory concordance in the TRACERx cohort (n = 213 specimens from 64 patients). The observed concordance rate (fraction of region pairs within a patient sharing the same trajectory assignment; red dashed line) is compared to a null distribution generated by random permutation of trajectory labels across regions. p value reflects the empirical permutation test. (E–F) Whole-genome instability index (WGII) (D) and number of driver alterations (E) stratified by TS. (G) Patient-level distribution of Transcriptomic Stage (TS) by the published TRACERx Renal evolutionary subtype (EvoType^41^). Per-patient TS was assigned as the latest call observed across that patient’s sampled regions (Late if any region was Late; otherwise Intermediate if any region was Intermediate; otherwise Early; otherwise Other), reflecting that progression in any sampled region marks the tumor as having reached that stage. Patient level EvoType labels were published for this dataset. Numbers above each bar indicate the number of patients in each EvoType group (n = 79 total).

**Figure S14.**
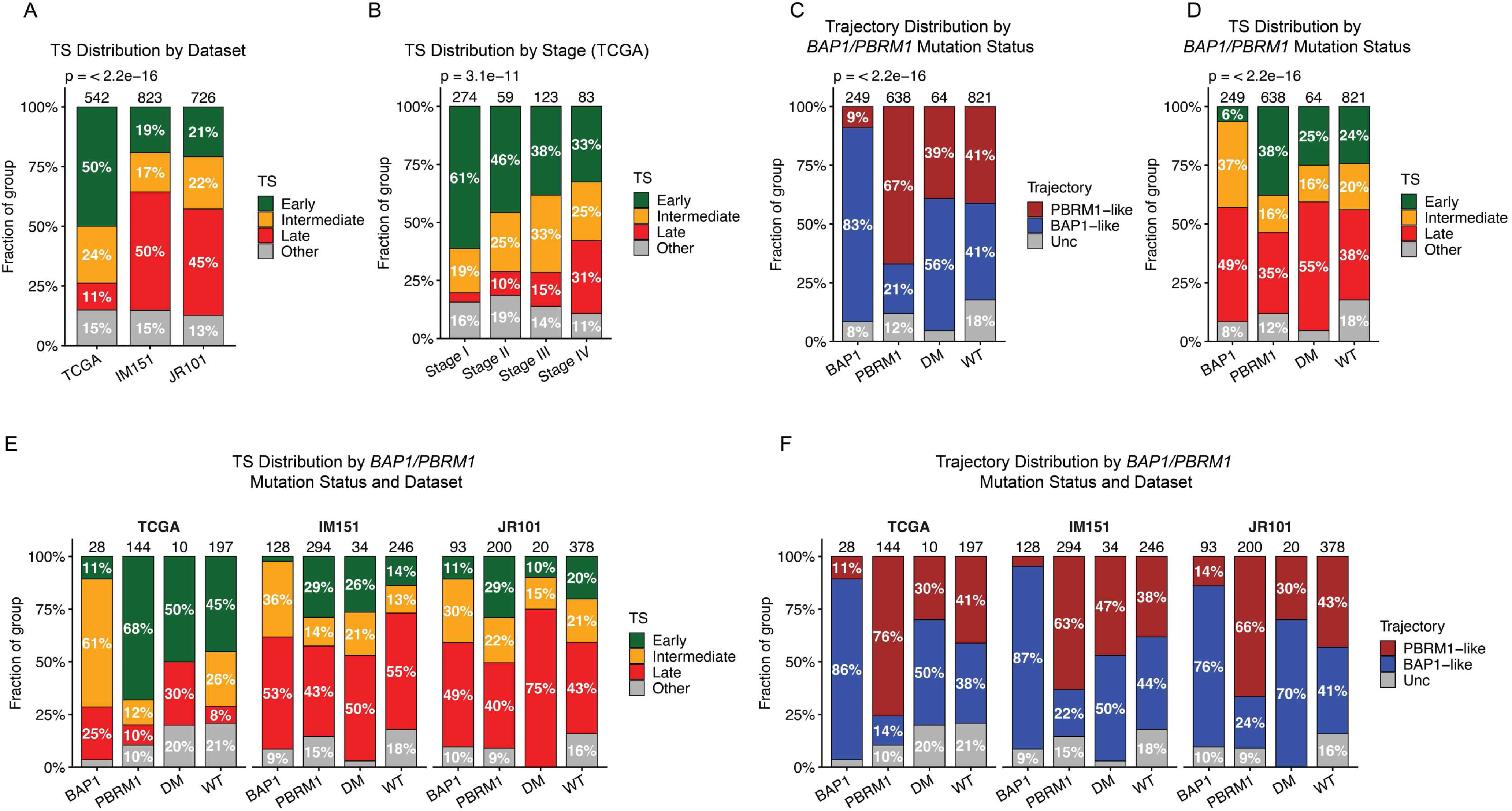
Distribution of transcriptomic stage differs by clinical setting (Related to Figure 6). (A) Distribution of TS stratified by dataset. (B) For the TCGA dataset, distribution of transcriptomic stage stratified by clinical stage. (C) Distribution of trajectory (*BAP1*-like, *PBRM1*-like, Unclassified) by *BAP1/PBRM1* genotype across all three cohorts combined. *BAP1*-mutant, *PBRM1*-mutant, *BAP1/PBRM1* double-mutant (DM), and *BAP1/PBRM1* wild-type (WT) groups are shown. (D) Distribution of TS by *BAP1/PBRM1* genotype, pooled across the three cohorts. (E) Distribution of TS by *BAP1/PBRM1* genotype, stratified by dataset (TCGA, IM151, JR101).

**Figure S15.**
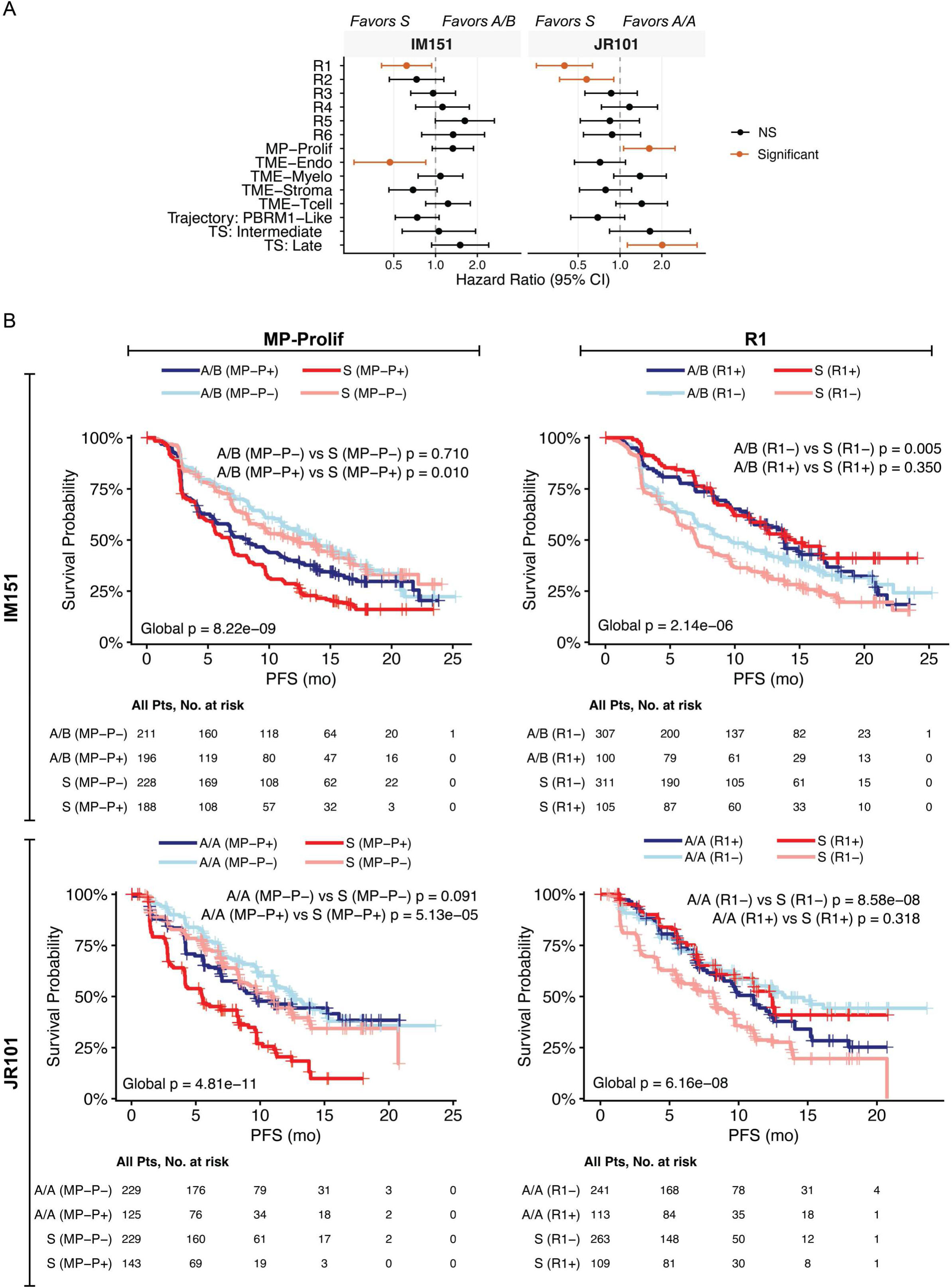
R1 and MP-Prolif utilization predict differential benefit from sunitinib (Related to Figure 6). (A) Forest plot of the treatment arm by CTP utilization interaction term from Cox proportional hazards models for the IM151 (left) and JR101 (right) cohorts, testing the association between each CTP (utilized vs not utilized) and outcome by treatment arm. Points denote hazard ratios (HRs) and bars denote 95% confidence intervals. (B) Kaplan–Meier analyses stratified by CTP utilization and treatment arm.

