## Supplementary Tables 1, 4-8 for "Clear cell renal cell carcinoma consensus transcriptomic programs reveal converging trajectories towards aggressive disease"

Table S1. Cohort Characteristics

|  | | **Dataset** | | |
| --- | --- | --- | --- | --- |
|  | **N = 2,163** | **IM151** N = 823 | **JR101** N = 726 | **TCGA** N = 614 |
| **Tissue Type** |  |  |  |  |
| *Normal* | 72 (3.3%) | 0 (0.0%) | 0 (0.0%) | 72 (11.7%) |
| *Tumor* | 2,091 (96.7%) | 823 (100.0%) | 726 (100.0%) | 542 (88.3%) |
| **DNAseq Available** |  |  |  |  |
| *No* | 391 (18.1%) | 121 (14.7%) | 35 (4.8%) | 235 (38.3%) |
| *Yes* | 1,772 (81.9%) | 702 (85.3%) | 691 (95.2%) | 379 (61.7%) |
| ***TFEB/TFE3* Fusion Annotation** |  |  |  |  |
| *No* | 798 (36.9%) | 0 (0.0%) | 726 (100.0%) | 72 (11.7%) |
| *Yes* | 1,365 (63.1%) | 823 (100.0%) | 0 (0.0%) | 542 (88.3%) |
| ***CDKN2A* Deletion Annotation** |  |  |  |  |
| *No* | 935 (43.2%) | 121 (14.7%) | 726 (100.0%) | 88 (14.3%) |
| *Yes* | 1,228 (56.8%) | 702 (85.3%) | 0 (0.0%) | 526 (85.7%) |
| **Sex** |  |  |  |  |
| *Female* | 595 (27.5%) | 229 (27.8%) | 178 (24.5%) | 188 (30.6%) |
| *Male* | 1,496 (69.2%) | 594 (72.2%) | 548 (75.5%) | 354 (57.7%) |
| *N/A* | 72 (3.3%) | 0 (0.0%) | 0 (0.0%) | 72 (11.7%) |
| **Nuclear Grade** |  |  |  |  |
| *G1* | 14 (0.6%) | 0 (0.0%) | 0 (0.0%) | 14 (2.3%) |
| *G2* | 237 (11.0%) | 0 (0.0%) | 0 (0.0%) | 237 (38.6%) |
| *G3* | 207 (9.6%) | 0 (0.0%) | 0 (0.0%) | 207 (33.7%) |
| *G4* | 76 (3.5%) | 0 (0.0%) | 0 (0.0%) | 76 (12.4%) |
| *N/A* | 1,629 (75.3%) | 823 (100.0%) | 726 (100.0%) | 80 (13.0%) |
| **Clinical Stage^1^** |  |  |  |  |
| *Stage I* | 274 (12.7%) | 0 (0.0%) | 0 (0.0%) | 274 (44.6%) |
| *Stage II* | 59 (2.7%) | 0 (0.0%) | 0 (0.0%) | 59 (9.6%) |
| *Stage III* | 123 (5.7%) | 0 (0.0%) | 0 (0.0%) | 123 (20.0%) |
| *Stage IV* | 1,632 (75.5%) | 823 (100.0%) | 726 (100.0%) | 83 (13.5%) |
| *N/A* | 75 (3.5%) | 0 (0.0%) | 0 (0.0%) | 75 (12.2%) |
| **IMDC Risk Group** |  |  |  |  |
| *Favorable* | 176 (8.1%) | 176 (21.4%) | 0 (0.0%) | 0 (0.0%) |
| *Intermediate* | 513 (23.7%) | 513 (62.3%) | 0 (0.0%) | 0 (0.0%) |
| *Poor* | 134 (6.2%) | 134 (16.3%) | 0 (0.0%) | 0 (0.0%) |
| *N/A* | 1,340 (62.0%) | 0 (0.0%) | 726 (100.0%) | 614 (100.0%) |
| **Treatment Arm** |  |  |  |  |
| *A/A* | 354 (16.4%) | 0 (0.0%) | 354 (48.8%) | 0 (0.0%) |
| *A/B* | 407 (18.8%) | 407 (49.5%) | 0 (0.0%) | 0 (0.0%) |
| *Sunitinib* | 788 (36.4%) | 416 (50.5%) | 372 (51.2%) | 0 (0.0%) |
| *N/A* | 614 (28.4%) | 0 (0.0%) | 0 (0.0%) | 614 (100.0%) |
| **OS Annotation** |  |  |  |  |
| *No* | 1,621 (74.9%) | 823 (100.0%) | 726 (100.0%) | 72 (11.7%) |
| *Yes* | 542 (25.1%) | 0 (0.0%) | 0 (0.0%) | 542 (88.3%) |
| **PFS Annotation** |  |  |  |  |
| *No* | 614 (28.4%) | 0 (0.0%) | 0 (0.0%) | 614 (100.0%) |
| *Yes* | 1,549 (71.6%) | 823 (100.0%) | 726 (100.0%) | 0 (0.0%) |
| **Median Duration of Follow Up Months (IQR)** |  | 16.6 (13.8-20.5) | 11.0 (7.0-15.2) | 55.1 (32.1-81.7) |

^1^Clinical stage was assigned at time of study enrollment in the IM151 and JR101 cohorts. In the TCGA it was defined at the time of nephrectomy. Abbreviations: A/A, avelumab and axitinib; A/B, atezolizumab and bevacizumab; IM151, IMmotion 151 Phase III study; IMDC, International Metastatic RCC Database Consortium Risk Score; IQR, Inter-quartile range; PFS, Progression free survival; JR101, Javelin Renal 101 Phase III study; OS, Overall survival; TCGA, The Cancer Genome Atlas Clear Cell Renal Cell Carcinoma Study.

Table S4. Driver mutation frequency in validation cohorts

| **Gene** | **CheckMate-025/010/009** | **TRACERx Renal** |
| --- | --- | --- |
| N annotated | 217 | 230 |
| *VHL* | 80 (36.9%) | 189 (82.2%) |
| *BAP1* | 26 (12.0%) | 41 (17.8%) |
| *PBRM1* | 70 (32.3%) | 108 (47.0%) |
| *TSC1* | 5 (2.3%) | 19 (8.3%) |
| *PTEN* | 13 (6.0%) | 12 (5.2%) |
| *TP53* | 3 (1.4%) | 8 (3.5%) |
| *KDM5C* | 26 (12.0%) | 42 (18.3%) |

Table S5. Multivariate analysis of TCGA dataset

| **Characteristic** | **N** | **HR** | **95% CI** | **p-value** |
| --- | --- | --- | --- | --- |
| **Stage** |  |  |  | <0.001 |
| *Stage I* | 270 | — | — |  |
| *Stage II* | 56 | 0.98 | 0.52, 1.82 |  |
| *Stage III* | 122 | 1.90 | 1.24, 2.92 |  |
| *Stage IV* | 83 | 4.27 | 2.75, 6.62 |  |
| **Nuclear Grade** |  |  |  | 0.077 |
| *G1/G2* | 249 | — | — |  |
| *G3* | 207 | 1.40 | 0.95, 2.06 |  |
| *G4* | 75 | 1.71 | 1.06, 2.76 |  |
| **Transcriptomic Stage** |  |  |  | <0.001 |
| *Early* | 268 | — | — |  |
| *Intermediate* | 128 | 1.75 | 1.17, 2.62 |  |
| *Late* | 60 | 2.42 | 1.56, 3.76 |  |
| *Other* | 75 | 1.65 | 1.03, 2.63 |  |
| Multivariate analysis of the TCGA dataset. Overall survival was modeled as a function of stage, grade, and transcriptomic stage. Analysis only includes patients with non-missing Stage or Nuclear Grade (n = 522). Abbreviations: CI, Confidence Interval; HR, Hazard Ratio. | | | | |

Table S6. Multivariate analysis of IM151 dataset

| **Characteristic** | **N** | **HR** | **95% CI** | **p-value** |
| --- | --- | --- | --- | --- |
| **Treatment Arm** |  |  |  | 0.034 |
| *S* | 416 | — | — |  |
| *A/B* | 407 | 0.83 | 0.70, 0.99 |  |
| **IMDC Risk Score** |  |  |  | <0.001 |
| *Favorable* | 176 | — | — |  |
| *Intermediate* | 513 | 1.46 | 1.15, 1.85 |  |
| *Poor* | 134 | 3.26 | 2.45, 4.34 |  |
| **Transcriptomic Stage** |  |  |  | <0.001 |
| *Early* | 157 | — | — |  |
| *Intermediate* | 136 | 1.12 | 0.83, 1.52 |  |
| *Late* | 408 | 1.57 | 1.23, 1.99 |  |
| *Other* | 122 | 0.90 | 0.65, 1.25 |  |
| Multivariable Cox proportional hazards analysis of progression-free survival in the IMmotion151 dataset, modeling progression-free survival as a function of treatment arm, IMDC risk score, and transcriptomic stage. Abbreviations: CI, Confidence Interval; HR, Hazard Ratio. | | | | |

Table S7. MVA of JR101 dataset

| **Characteristic** | **N** | **HR** | **95% CI** | **p-value** |
| --- | --- | --- | --- | --- |
| **Treatment Arm** |  |  |  | <0.001 |
| *S* | 372 | — | — |  |
| *A/A* | 354 | 0.66 | 0.54, 0.82 |  |
| **Transcriptomic Stage** |  |  |  | 0.002 |
| *Early* | 151 | — | — |  |
| *Intermediate* | 159 | 1.03 | 0.74, 1.44 |  |
| *Late* | 324 | 1.54 | 1.16, 2.04 |  |
| *Other* | 92 | 1.00 | 0.68, 1.48 |  |
| Multivariable Cox proportional hazards analysis of progression-free survival in the JAVELIN Renal 101 dataset, modeling progression-free survival as a function of treatment arm and transcriptomic stage. Abbreviations: CI, Confidence Interval; HR, Hazard Ratio. | | | | |

Table S8. MVA of the TCGA including BAP1/PBRM1 classifier

| **Characteristic** | **N** | **HR** | **95% CI** | **p-value** |
| --- | --- | --- | --- | --- |
| **Stage** |  |  |  | <0.001 |
| *Stage I* | 201 | — | — |  |
| *Stage II* | 42 | 0.71 | 0.29, 1.75 |  |
| *Stage III* | 77 | 1.46 | 0.81, 2.64 |  |
| *Stage IV* | 50 | 4.10 | 2.28, 7.35 |  |
| **Nuclear Grade** |  |  |  | 0.370 |
| *G1/G2* | 182 | — | — |  |
| *G3* | 140 | 1.26 | 0.74, 2.14 |  |
| *G4* | 48 | 1.59 | 0.83, 3.02 |  |
| ***BAP1/PBRM1* Status** |  |  |  | 0.848 |
| *WT* | 190 | — | — |  |
| *PBRM1* | 142 | 1.24 | 0.77, 2.01 |  |
| *BAP1* | 28 | 1.06 | 0.55, 2.03 |  |
| *DM* | 10 | 1.01 | 0.35, 2.92 |  |
| **Transcriptomic Stage** |  |  |  | 0.001 |
| *Early* | 192 | — | — |  |
| *Intermediate* | 84 | 2.28 | 1.25, 4.17 |  |
| *Late* | 39 | 3.50 | 1.89, 6.49 |  |
| *Other* | 55 | 1.53 | 0.80, 2.96 |  |
| Multivariable Cox proportional hazards analysis of overall survival in the TCGA dataset, modeling overall survival as a function of tumor stage, tumor grade, *BAP1/PBRM1* status, and transcriptomic stage. Analysis included only samples with Stage, Nuclear Grade and *BAP1/PBRM1* annotations (n = 370). Abbreviations: CI, Confidence Interval; HR, Hazard Ratio. | | | | |
